# An epi-evolutionary model to predict spore-producing pathogens adaptation to quantitative resistance in heterogeneous environments

**DOI:** 10.1101/423467

**Authors:** Frédéric Fabre, Jean-Baptiste Burie, Arnaud Ducrot, Sébastien Lion, Quentin Richard, Ramsès Djidjou-Demasse

**Affiliations:** INRAE, Bordeaux Sciences Agro, UMR SAVE, Villenave d’Ornon F-33882, France; CNRS, IMB, UMR 5251, Talence F-33400, France; Univ. Bordeaux, IMB, UMR 5251, Talence F-33400, France; Univ. Normandie, UNIHAVRE, LMAH, FR-CNRS-3335, ISCN, 76600 Le Havre, France; CEFE, CNRS, Univ. Montpellier, Univ. Montpellier 3 Paul-Valéry, EPHE, IRD, Montpellier F-34293, France; MIVEGEC, Univ. Montpellier, IRD, CNRS, Montpellier, France

**Keywords:** Basic reproduction number, Resistance durability, Adaptive dynamics, Spore-producing pathogens, Quantitative resistance

## Abstract

We model the evolutionary epidemiology of spore-producing plant pathogens in heterogeneous environments sown with several cultivars carrying quantitative resistances. The model explicitly tracks the infection-age structure and genetic composition of the pathogen population. Each strain is characterized by pathogenicity traits describing its infection efficiency and a time-varying sporulation curve taking into account lesion ageing. We first derive a general expression of the basic reproduction number ℛ_0_ for fungal pathogens in heterogeneous environments. We show that evolutionary attractors of the model coincide with local maxima of ℛ_0_ only if the infection efficiency is the same on all host types. We then study how three basic resistance characteristics (pathogenicity trait targeted, resistance effectiveness, and adaptation cost) in interaction with the deployment strategy (proportion of fields sown with a resistant cultivar) (i) lead to pathogen diversification at equilibrium and (ii) shape the transient dynamics from evolutionary and epidemiological perspectives. We show that quantitative resistance impacting only the sporulation curve will always lead to a monomorphic population, while dimorphism (*i*.*e*. pathogen diversification) can occur with resistance altering infection efficiency, notably with high adaptation cost and proportion of R cultivar. Accordingly, the choice of quantitative resistance genes operated by plant breeders is a driver of pathogen diversification. From an evolutionary perspective, the emergence time of the evolutionary attractor best adapted to the R cultivar tends to be shorter when the resistance impacts infection efficiency than when it impacts sporulation. In contrast, from an epidemiological perspective, the epidemiological control is always higher when the resistance impacts infection efficiency. This highlights the difficulty of defining deployment strategies of quantitative resistance maximising at the same time epidemiological and evolutionary outcomes.

## 1 Introduction

Resistance to parasites, i.e., the capacity of a host to decrease its parasite development (Raberg et *al*., 2009), is a widespread defense mechanism in plants. Qualitative resistance usually confers disease immunity in such a way that parasites exhibit a discrete distribution of their disease phenotype (“cause disease” versus “do not cause disease”) on a resistant plant (McDonald & Linde, 2002). Quantitative resistance leads to a reduction in disease severity (Poland et *al*., 2009; St. Clair, 2010) in such a way that parasites exhibit a continuous distribution of their pathogenicity (McDonald & Linde, 2002; St. Clair, 2010; Lannou, 2012). Pathogenicity, also termed aggressiveness, can be estimated in laboratory experiments through the measure of a small number of pathogenicity traits (Lannou, 2012) expressed during the basic steps of the host-pathogen interaction. Quantitative resistance has attracted attention in plant breeding for pathogen control in low-input cropping systems, in particular due to their supposed higher durability compared with qualitative resistance (Niks et *al*., 2015). However, plant pathogens also adapt to quantitative resistance (see Pilet-Nayel et *al*. (2017) for a review). The resulting gradual “erosion” of resistance effectiveness (McDonald & Linde, 2002) corresponds, from the pathogen side, to a gradual increase in pathogenicity.

Historically, theoretical studies investigating how the deployment of quantitative resistance in agrosystems impacts pathogen aggressiveness have relied on adaptive dynamics (*e*.*g*. Gudelj et *al*. (2004a,b); van den Bosch et *al*. (2006, 2007); van den Berg et *al*. (2014)). Adaptive dynamics (Geritz et *al*., 1997, 1998; Dieckmann, 2002) supposes that epidemiological and evolutionary processes unfold on at different time scales. It essentially focuses on long-term predictions for endemic diseases. Fewer studies (Iacono et *al*., 2012; Bourget *al*., 2015; Rimbaud et *al*., 2018) address the fundamental short- and long-term objectives of sustainable management of plant diseases (Zhan et *al*., 2015; Rimbaud et *al*., 2018): the short-term goal focuses on the reduction of disease incidence, whereas the longer-term objective is to reduce pathogen adaptation to resistant cultivars. Evolutionary epidemiology analysis is well-suited for this purpose (Day & Proulx, 2004). Essentially inspired by quantitative genetics, it accounts for the interplay between epidemiological and evolutionary dynamics on the same time scale. As such it can be used to monitor the simultaneous dynamics of epidemics and the evolution of any set of pathogen life-history trait of interest. It can also take into account the heterogeneity of host populations resulting, for example, from differences in the genetic, physiological, or ecological states of individuals (Day & Gandon, 2007). This is typically the case with field mixtures, where several cultivars are cultivated in the same field, and with landscape mosaics, with cultivars cultivated in different fields (Rimbaud et *al*., 2021).

In this article, we follow this approach and study the evolutionary epidemiology of spore-producing pathogens in heterogeneous agricultural environments. Plant fungal pathogens (*sensu lato, i*.*e*., including Oomycetes) are typical spore-producing pathogens responsible for nearly one third of emerging plant diseases (Anderson et *al*., 2004). Among their common features, spore production is usually a function of the time since infection as a result of lesion ageing (van den Bosch et *al*., 1988; Sache et *al*., 1997; Kolnaar & Bosch, 2011; Segarra et *al*., 2001). We first formulate a general model that explicitly tracks the infection-age structure and genetic composition of the pathogen population. Mathematically, the model is an extension to heterogeneous plant populations of an integro-differential model introduced by Djidjou-Demasse et *al*. (2017). We then investigate how the deployment of quantitative resistances altering different pathogenicity traits impacts the pathogen population structure at equilibrium. This question is addressed by highlighting the links between the frameworks of evolutionary epidemiology and adaptive dynamics. We characterize the evolutionary attractors of the coupled epidemiological evolutionary dynamics while emphasising the differences between the cornerstone concepts of ℛ_0_ in epidemiology (Diekmann et *al*., 1990; van den Driessche & Watmough, 2008) and invasion fitness in evolution (Dieckmann, 2002; Diekmann et *al*., 2005; Geritz et *al*., 1998; Metz et *al*., 1996; Nowak & Sigmund, 2004). Finally, we investigate how the deployment of quantitative resistances impacts the transient behavior of the coupled epi-evolutionary dynamics, both at epidemiological (disease incidence) and evolutionary levels (resistance durability).

## 2 An epi-evolutionary model for spore-producing pathogens

Suppose we cultivate a field with two host types that differ by their quantitative level of resistance to a pathogen (say a resistant and a susceptible cultivar). How can we model the joint epidemiological and evolutionary dynamics of the host-pathogen interaction? In this section, we first formulate a general model for the dynamics of spore-producing plant pathogens in structured populations, which we then apply to study specific scenarios later on.

### 2.1 Host and pathogen populations

We consider a spore-producing plant pathogen infecting a heterogeneous host population with *N*_*c*_ host classes. In our study, the host classes are assumed to represent different plant cultivars, but more generally host heterogeneity may represent different host developmental or physiological states or different habitats. At a given time *t*, hosts in class *k* can be either healthy (*H*_*k*_(*t*)) or infected. Note that, in keeping with the biology of fungal pathogens, we do not track individual plants, but rather leaf area densities (leaf surface area per m^2^). The leaf surface is viewed as a set of individual patches corresponding to a restricted host surface area that can be colonized by a single pathogen individual (*i*.*e*. no coinfection is possible). Thus, only indirect competition between pathogen strains for a shared resource is considered. Spores produced by all infectious tissues are assumed to mix uniformly in the air. They constitute a well-mixed pool of spores landing on any host class according to the law of mass action. Thus, the probability of contact between a spore and host *k* is proportional to the total healthy leaf surface area of this host.

The parasite population is structured by a continuous phenotype *x* and by the age of infection *a*, so that *I*_*k*_(*a, t, x*) represents the density of infected tissue in host class *k* at time *t* that were infected *t* −*a* time units ago by a pathogen strain *x*. The density of the airborne pool of spores with phenotype *x* at time *t* is denoted by *A*(*t, x*).

Both the phenotype and age of infection affect on each host class two pathogenicity traits summarising the basic steps of the disease infection cycle: (i) the infection efficiency *β*_*k*_(*x*), *i*.*e*., the probability that a spore deposited on a receptive host surface produces a lesion and (ii) the sporulation curve *r*_*k*_(*a, x*). In line with the biology of plant fungi (van den Bosch et *al*., 1988; Sache et *al*., 1997; Kolnaar & Bosch, 2011; Segarra et *al*., 2001), a gamma sporulation curve defined by four parameters is assumed: (i) the latent period *τ*_*k*_(*x*), (ii) the total number of spores *p*_*k*_(*x*) produced by a lesion during its whole infectious period and (iii-iv) the rate and shape parameters of the gamma distribution (respectively denoted *λ*_*k*_(*x*) and *n*_*k*_(*x*)). These hypotheses lead to the following sporulation function :

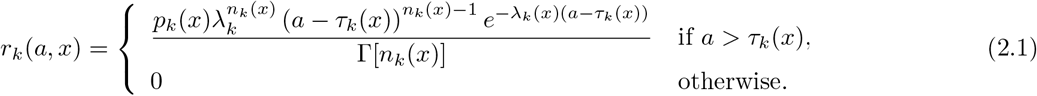

where Γ is the Gamma function. With this formalism, the duration of the infectious period can be estimated (based on a Gaussian approximation) as 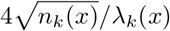.

### 2.2 Epi-evolutionary dynamics

With these assumptions, we can derive the following integro-differential equations which describe the epidemiological and evolutionary dynamics of the host and pathogen populations (see Figure 1C for a schematic representation of the model with 2 hosts):

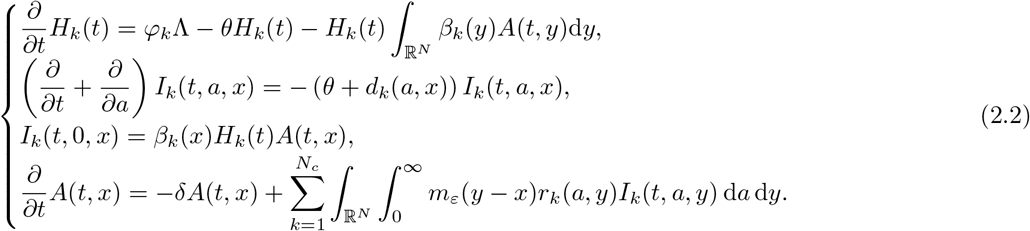

Healthy hosts are produced at rate Λ and *φ*_*k*_ is the proportion of the host *k* at planting in the environment. Healthy hosts can become infected by airborne spores. The total force of infection on a host 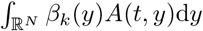, where ℛ^*N*^ is the phenotype space of dimension *N*. Airborne spores produced by infected hosts become nonviable at rate *δ*. Healthy hosts die at rate *θ* (regardless of their class) and infected hosts at rate *θ* + *d*_*k*_(*a, x*), where *d*_*k*_(*a, x*) is the disease-induced mortality. Hosts infected by strain *y* produce airborne spores with phenotype *x* at rate *m*_*ε*_(*y − x*)*r*_*k*_(*a, y*), where *m*_*ε*_(*y − x*) is the probability of mutation from phenotype *y* to phenotype *x*. Thus, mutations randomly displace strains into the phenotype space at each infection cycle (*i*.*e*., generation) according to a mutation kernel *m*_*ε*_. A centered multivariate Gaussian distribution with standard deviation *ε* is typically used thereafter. However, other mutation kernels can be used as soon as they satisfy some general properties (Appendix B).

**Figure 1:**
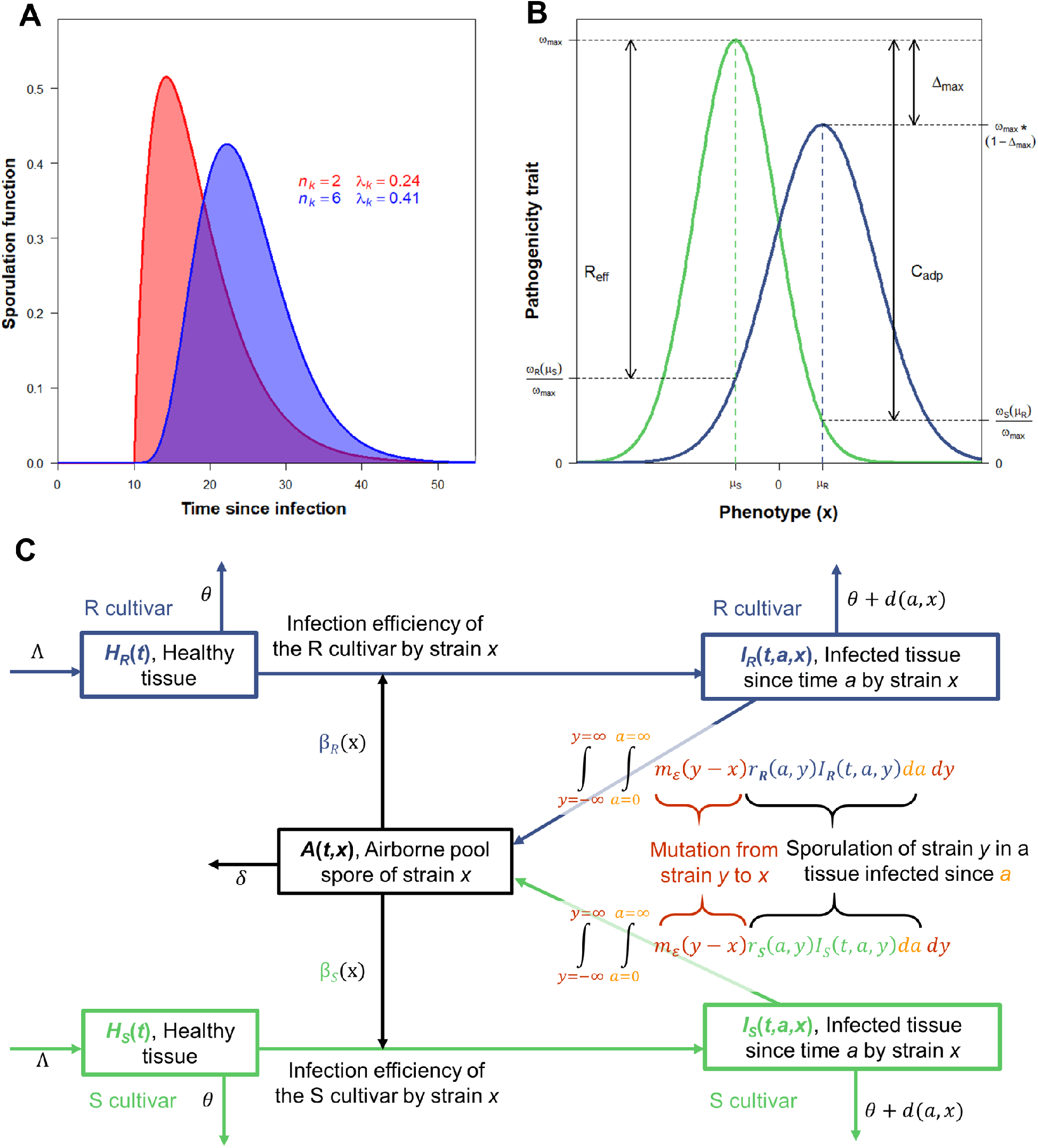
**A** Shapes of gamma sporulation function *r*_*k*_(*a, x*) with latent period *τ*_*k*_ = 10 and total spore production during the whole infectious period *p*_*k*_ = 5.94 as a function of *n*_*k*_ and *λ*_*k*_. The infectious period is =∼ 24 days for both functions. **B** Phenotypic landscapes of a pathogenicity trait *ω* on a susceptible (S) and on a resistant (R) cultivar. Trait values are described by unnormalized Gaussian functions. The optimal parasite phenotypes *µ*_*S*_ and *µ*_*R*_ have maximum traits *ω*_max_ and *ω*_max_(1 − Δ_*max*_) on the S and R cultivars, respectively. The relative effectiveness of resistance *R*_eff_ = (*ω*_*S*_(*µ*_*S*_) − *ω*_*R*_(*µ*_*S*_)) */ω*_max_ corresponds to the trait difference between the cultivars of the phenotype *µ*_*S*_. The relative cost of adaptation *C*_adp_ = (*ω*_*S*_(*µ*_*S*_) − *ω*_*S*_(*µ*_*R*_)) */ω*_max_ corresponds to the trait difference between the phenotypes *µ*_*S*_ and *µ*_*R*_ on the S cultivar. Here *R*_*eff*_ = 0.8, *C*_adp_ = 0.9 and Δ_*max*_ = 0.2. **C** Schematic representation of the model with 2 hosts (a susceptible (S) and a resistant (R) cultivar) and an unidimensionnal mutation kernel (*i*.*e. N*_*c*_ = 2 and *N* = 1). All state variables and parameters are listed in Table 1.

In the simplest scenarios, the phenotype space can be one-dimensional (*i*.*e. N* = 1), but we can also consider a multi-dimensional phenotype space (*i*.*e. N* ≥ 2). We refer the reader to Table 1 for a list of the variables and parameters of the model, and to Appendix C for a discussion of how to recover several simpler models in the literature from System (2.2).

**Table 1:** Main notations, state variables and parameters of the model. The general model is structured by age of infection *a*, pathogen strain *x* and host class *k*. Some of its parameters are functions of these structuring classes.

| Category | Description | Unit |
| --- | --- | --- |
| Notations |  |  |
| $k \in 1, 2, \dots, N_c$ | Index on host class ($N_c \geq 2$ class are considered) | |
| $\mathbb{R}^N$ | Phenotype space of dimension $N$ | |
| $x \in \mathbb{R}^N$ | Label of the pathogen strain | |
| $a$ | Time since infection | |
| States variables |  |  |
| $H_k(t)$ | Density of healthy tissue of host $k$ at time $t$ | |
| $I_k(t, a, x)$ | Density of infected tissue of host $k$ at time $t$ infected since time $a$ by strain $x$ | |
| $A(t, x)$ | Density of airborne pool of pathogen spores of strain $x$ at time $t$ | |
| Parameters |  |  |
| $\Lambda$ | Birth rate of healthy host tissue | HTD · Tu$^{-1}$ |
| $\theta$ | Death rate of host tissue | Tu$^{-1}$ |
| $\delta$ | Rate at which spore becomes nonviable | Tu$^{-1}$ |
| $d_k(a, x)$ | Disease induced mortality of host $k$ infected since $a$ by strain $x$ | Tu$^{-1}$ |
| $\beta_k(x)$ | Infection efficiency of pathogen strain $x$ on host $k$ | SD$^{-1} \cdot$ Tu$^{-1}$ |
| $p_k(x)$ | Total number of spores produced by a lesion caused by strain $x$ on host $k$ | SD · HTD$^{-1}$ |
| $\tau_k(x)$ | Latent period of pathogen strain $x$ on host $k$ | Tu |
| $\lambda_k(x)$ | Rate of the gamma sporulation curve shape for strain $x$ on host $k$ | unitless |
| $n_k(x)$ | Shape of the gamma sporulation curve shape for strain $x$ on host $k$ | unitless |
| $\varphi_k$ | Proportion of host $k$ at planting | unitless |
| $\varepsilon$ | Standard deviation of the Gaussian mutation kernel | unitless |

Tu= time unit; SD= spores density; HTD=host tissue density

## 3 Basic reproduction number and invasion fitness

The model (2.2) allows us to track the coupled epidemiological and evolutionary dynamics of the host-pathogen interaction. We can use a numerical integration of this model to determine the transient dynamics towards potential evolutionary attractors. To make further analytical progress, we can consider two limiting cases of this process. First, when the population is initially uninfected, whether a single pathogen strain spreads or not in the population can be determined by calculating the basic reproduction number of this strain. Second, the long-term evolutionary endpoints can be determined by calculating the invasion fitness of a rare mutant in a resident population at equilibrium, following the standard adaptive dynamics methodology.

### 3.1 Basic reproduction number: Invasion in an uninfected population

The basic reproduction number, usually denoted as ℛ_0_, is defined as the total number of infections arising from one newly infected individual introduced into a healthy (disease-free) host population (Diekmann et *al*., 1990; Anderson, 1991). It can typically be used to study the spread of a pathogen strain *x* in an uninfected host population. In an environment with *N*_*c*_ host classes where the pathogen propagules all pass through a common pool in the compartment *A* as in model (2.2), ℛ_0_ is the sum of the pathogen’s reproductive numbers for each host class (Rueffler & Metz, 2013). A pathogen with phenotype *x* will spread if ℛ_0_(*x*) *>* 1, with

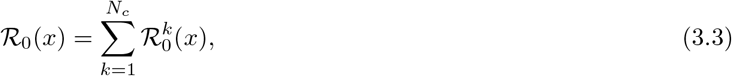

where the quantity 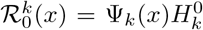 is the basic reproduction number of a pathogen with phenotype *x* in host *k*. This expression depends on the disease-free equilibrium density of healthy hosts 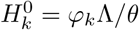 and Ψ_*k*_(*x*) the reproductive value of a pathogen with phenotype *x* landing on host *k* (*i*.*e*. the fitness function, Appendix D). It is given by

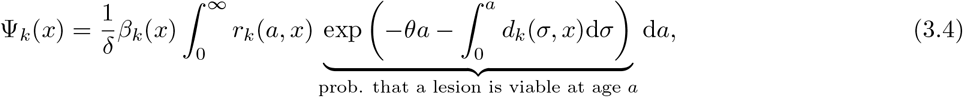

In this expression, multiplying the probability that a lesion is viable at age *a* by *r*_*k*_(*a, x*) and integrating over all infection ages *a* gives the total number of spores effectively produced by a lesion during its lifetime. It differs from *p*_*k*_(*x*) which is the number of spores potentially produced if the lesion remains viable during the whole infectious period. A close expression with a simpler model (single host type and no disease-induced mortality) was derived by Gilchrist et *al*. (2006).

Assuming that *d*_*k*_ does not depend on the age of infection and a gamma sporulation function (2.1), we obtain

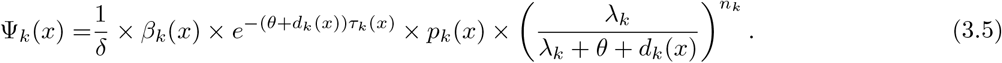

Furthermore, if *r*_*k*_ is a constant (*i*.*e. r*_*k*_(*a, x*) = *p*_*k*_(*x*) and *d*_*k*_(*a, x*) = *d*_*k*_(*x*) for all *a*), we recover the classical expression of ℛ_0_ for SIR models (Day, 2002) with

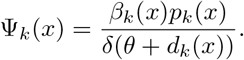

### 3.2 Invasion fitness and long-term evolutionary attractors

Once a pathogen strain *x* has reached an endemic equilibrium, the invasion of a new mutant strain can be studied using adaptive dynamics methodology. A rare mutant strain with phenotype *y* will invade a resident pathogen population with phenotype *x* if its invasion fitness *f*_*x*_(*y*) *>* 0. Potential evolutionary endpoints can be identified as the solutions of *f*_*x*_(*y*) = 0. The sign of this two-dimensional function is classically visualized using Pairwise Invasibility Plot (PIP) (Dieckmann, 2002; Diekmann et *al*., 2005; Geritz et *al*., 1997, 1998; Metz et *al*., 1996; Nowak & Sigmund, 2004). In our model, the existence of a common pool of pathogen propagules allows us to write (Appendix F) the invasion fitness *f*_*x*_(*y*) as

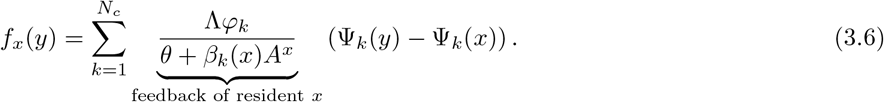

The environmental feedback of the resident strain *x* conditions the ability of a mutant strain *y* to invade the resident population. It depends on the conditions set out by the resident, in particular on the resources of host *k* already taken by *x*. When infection efficiencies do not differ between host classes (*i*.*e*., *β*_*k*_ = *β*, for all host classes *k*), this feedback term is unique and comes out of the sum, and using equation (3.3), equality (3.6) can be rewritten as

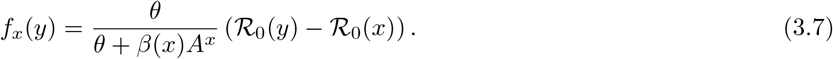

It follows that the model (2.2) admits an optimisation principle based on ℛ_0_ (Mylius & Diekmann, 1995; Metz et *al*., 2008; Rueffler & Metz, 2013; Lion & Metz, 2018; Gyllenberg & Service, 2011). Indeed, the sign of the invasion fitness *f*_*x*_(*y*) is given by the sign of the difference between ℛ_0_(*y*) and ℛ_0_(*x*) and thus, the evolutionary attractors of model (2.2) coincide with the local maxima of the ℛ_0_ as soon as host plants impact only sets of pathogenicity traits involved in the sporulation curves. Conversely, if infection efficiencies differ for at least two host classes the optimization principle does not apply. Accordingly, as soon as some plant hosts impact efficiency, the calculation of invasion fitness using equation (3.6) becomes necessary to characterize evolutionary attractors.

## 4 Case study: Deployment of quantitative resistances

As an application of our general model, we now consider two habitats corresponding to a susceptible (S) and a resistant (R) cultivar that differ by a single quantitative resistance trait. We assume that, after a long time of monoculture of the S cultivar, a fraction *ϕ* of the S cultivar is replaced by the R cultivar at *t* = 0. We consider two scenarios, depending on how the resistance trait impacts the life cycle of the pathogen. In the SP scenario, the resistance alters the total spore production only, while in the IE scenario, the resistance affects infection efficiency only. The analysis of the scenarios borrows analytical results from adaptive dynamics and simulation results from evolutionary epidemiology while highlighting the links between these frameworks.

### 4.1 scenarios simulations and model outputs

#### Phenotypic landscapes on the S and R cultivars

Simulating the model first requires specifying fitness functions describing pathogenicity trait values of any strain *x* on each cultivar (Figure 1B). For a trait *ω* (either the total spore production in the SP scenario, either the infection efficiency in the IE scenario), we used the unnormalized Gaussian function

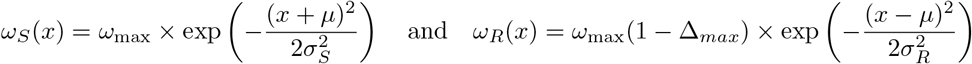

Without loss of generality, and to simplify the presentation, the optimal phenotypes are opposite for the S and R cultivars. Therefore, the optimal parasite phenotypes *µ*_*S*_ = −*µ* and *µ*_*R*_ = *µ* are respectively characterized by maximal trait values *ω*_max_ and *ω*_max_(1 − Δ_*max*_) on the S and R cultivars, where Δ_*max*_ is the relative difference of maximal traits between both cultivars. The inverse of the variances 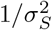 and 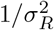 define the selectivity of each habitat. Rather than using these parameters, the phenotypic landscapes can be re-parameterized to fit the terminology used in plant pathology to describe quantitative resistances with the relative effectiveness of resistance *R*_eff_ and the relative cost of adaptation *C*_adp_ (Figure 1B). More precisely, for given *σ*_*S*_ *>* 0, *R*_eff_ and *C*_adp_ both in [0, 1] and Δ_*max*_ in [0, *R*_eff_[, we have

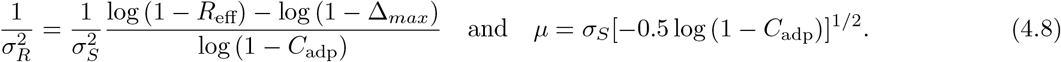

#### Parameter values and initial conditions

For each scenario, simulations were used to explore how the relative effectiveness of resistance *R*_eff_ (between 0.5 and 0.99), the relative cost of adaptation *C*_adp_ (between 0.3 and 0.99) and the deployment strategy *φ* (between 0.05 and 0.95) affect the coupled epidemiological and evolutionary dynamics. Note that, within the parameter space considered, the trade-off function that links *ω*_*S*_ and *ω*_*R*_ in both cultivars can be either concave, convex or sigmoidal (Figure S1). Similarly, the basic reproduction ratio ℛ_0_ can have either one global maximum, or one global and one local maximum while this global maximum can be closer to *µ*_*S*_ or to *µ*_*R*_ (Figure 4 line 3). These features strongly affect the dynamics. Simulations were initiated with a density of airborne spores *A*(0, *x*) at mutation-selection equilibrium in an environment where only the S cultivar was grown.

All model parameters and initial conditions are summarized in Table 2. Parameters for pathogenicity traits do not fit a particular pathogen species but rather typical biotrophic foliar fungal diseases, such as wheat rusts on susceptible cultivars. Following Rimbaud et *al*. (2018), we define an infection efficiency of 0.2 and set the duration of the latent and infectious periods to 10 and 24 days, respectively. The total spore production is set in order to obtain an ℛ_0_ of 30 in an environment with only the S cultivar (Mikaberidze et *al*., 2016).

**Table 2:**
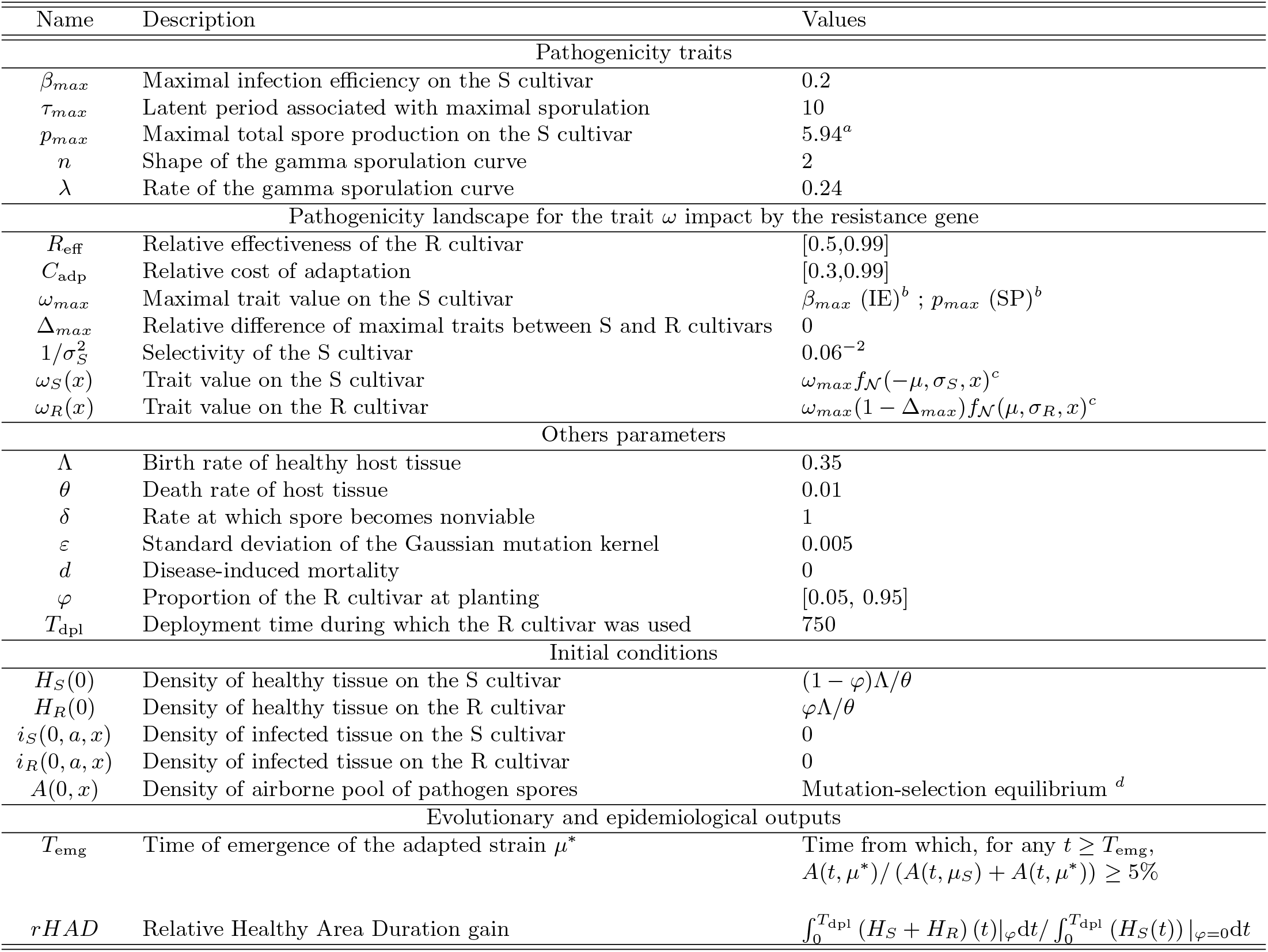
Initial conditions and parameters used for the simulations with the S and R cultivars. Two scenarios were considered: the R cultivar can either alter the Infection Efficiency of the pathogen (IE) or its total Spore Production (SP). Reference values given for each parameter are the one used in all simulations, unless stated otherwise. Values in brackets indicate the range of variation [minimum, maximum] used for the numerical exploration. ^*a*^ *p*_*max*_ is such that ℛ_0_(*µ*_*S*_) = 30 for *φ* = 0, leading to *p*_*max*_ = 30 *× θδ* exp(*θτ*_*max*_)(1 + *θ/λ*)^*n*^*/*(*β*_*max*_Λ).^*b*^ *ω*_*max*_ = *β*_*max*_ for the IE scenario and *ω*_*max*_ = *p*_*max*_ for the SP scenario. ^*c*^ The notations *f*_𝒩_ (*µ, σ, x*) = exp[−(*x* − *µ*)^2^*/*(2*σ*^2^)] stand for the unnormalized density function of the Gaussian distribution. The optimal phenotype value *µ* and the selectivity of the R cultivar 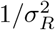 are calculated from equation (4.8). ^*d*^ The density of airborne pool of pathogen spores at mutation-selection equilibrium is determined by running the model during 3000 generations in an environment with only the S cultivar.

| Name | Description | Values |
| --- | --- | --- |
| Pathogenicity traits |  |  |
| $\beta_{max}$ | Maximal infection efficiency on the S cultivar | 0.2 |
| $\tau_{max}$ | Latent period associated with maximal sporulation | 10 |
| $p_{max}$ | Maximal total spore production on the S cultivar | $5.94^a$ |
| $n$ | Shape of the gamma sporulation curve | 2 |
| $\lambda$ | Rate of the gamma sporulation curve | 0.24 |
| Pathogenicity landscape for the trait $\omega$ impact by the resistance gene | | |
| $R_{eff}$ | Relative effectiveness of the R cultivar | [0.5,0.99] |
| $C_{adp}$ | Relative cost of adaptation | [0.3,0.99] |
| $\omega_{max}$ | Maximal trait value on the S cultivar | $\beta_{max}$ (IE)<sup>b</sup> ; $p_{max}$ (SP)<sup>b</sup> |
| $\Delta_{max}$ | Relative difference of maximal traits between S and R cultivars | 0 |
| $1/\sigma_S^2$ | Selectivity of the S cultivar | $0.06^{-2}$ |
| $\omega_S(x)$ | Trait value on the S cultivar | $\omega_{max} f_{\mathcal{N}}(-\mu, \sigma_S, x)^c$ |
| $\omega_R(x)$ | Trait value on the R cultivar | $\omega_{max}(1 - \Delta_{max}) f_{\mathcal{N}}(\mu, \sigma_R, x)^c$ |
| Others parameters |  |  |
| $\Lambda$ | Birth rate of healthy host tissue | 0.35 |
| $\theta$ | Death rate of host tissue | 0.01 |
| $\delta$ | Rate at which spore becomes nonviable | 1 |
| $\varepsilon$ | Standard deviation of the Gaussian mutation kernel | 0.005 |
| $d$ | Disease-induced mortality | 0 |
| $\varphi$ | Proportion of the R cultivar at planting | [0.05, 0.95] |
| $T_{dpl}$ | Deployment time during which the R cultivar was used | 750 |
| Initial conditions |  |  |
| $H_S(0)$ | Density of healthy tissue on the S cultivar | $(1 - \varphi)\Lambda/\theta$ |
| $H_R(0)$ | Density of healthy tissue on the R cultivar | $\varphi\Lambda/\theta$ |
| $i_S(0, a, x)$ | Density of infected tissue on the S cultivar | 0 |
| $i_R(0, a, x)$ | Density of infected tissue on the R cultivar | 0 |
| $A(0, x)$ | Density of airborne pool of pathogen spores | Mutation-selection equilibrium<sup>d</sup> |
| Evolutionary and epidemiological outputs |  |  |
| $T_{emg}$ | Time of emergence of the adapted strain $\mu^*$ | Time from which, for any $t \geq T_{emg}$,<br>$A(t, \mu^*) / (A(t, \mu_S) + A(t, \mu^*)) \geq 5\%$ |
| $rHAD$ | Relative Healthy Area Duration gain | $\int_0^{T_{dpl}} (H_S + H_R)(t) \vert _{\varphi} dt / \int_0^{T_{dpl}} (H_S(t)) \vert _{\varphi=0} dt$ |

#### Evolutionary and epidemiological outputs

The evolutionary output considered was the time of emergence *T*_emg_ of the adapted strain *µ*\*. It corresponds to the time from which its proportion in the airborne pool of spore compared to *µ*_*S*_ remains ≥ 0.05 (Table 2). If the equilibrium is monomorphic, the adapted strain *µ*\* is this unique evolutionary attractor. If the equilibrium is polymorphic, the adapted strain *µ*\* considered is the one closer to *µ*_*R*_. The epidemiological control provided by the deployment of the R cultivar was assessed with the relative Healthy Area Duration (rHAD) gain, a variable considered as a proxy for crop yield (Iacono et *al*., 2012; Rimbaud et *al*., 2018). It was calculated over 750 generations by assuming that a R cultivar is deployed over 50 years and the pathogen realized 15 generations each year (Table 2).

### 4.2 Evolutionary outcomes

In the SP scenario, the pathogenicity trait targeted by the quantitative resistance is only the total spore production. As the ℛ_0_ optimization principle holds, the population always becomes monomorphic around an evolutionary attractor *µ*\* corresponding to the global maximum of the ℛ_0_ function. It happens regardless of the shape of ℛ_0_, whether ℛ_0_ has a unique maximum (Figure 2 line 1) or it has an additional local maximum (Figure 2 line 2).

**Figure 2:**
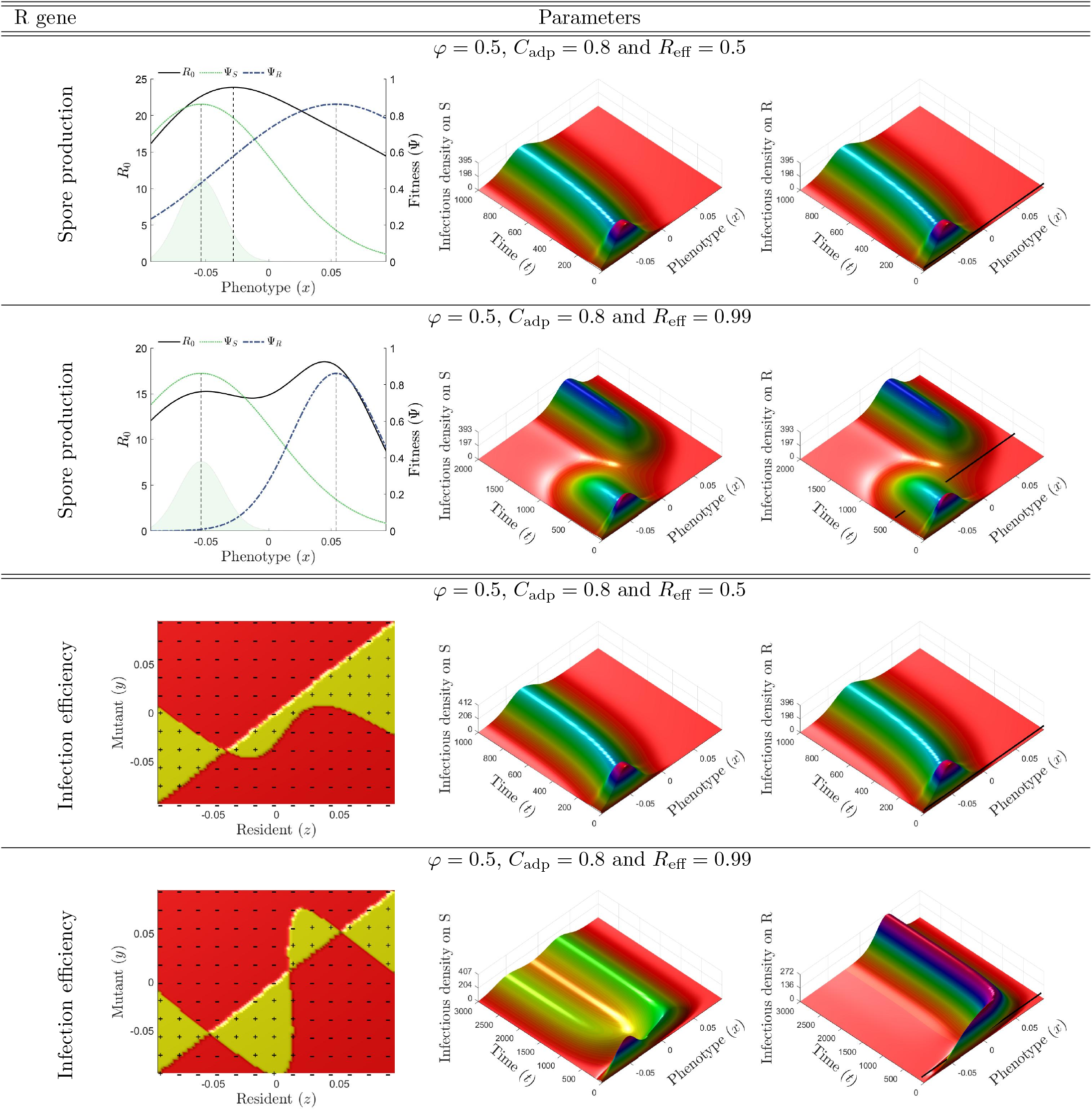
Evolutionary epidemiological dynamics. **Line 1**. The resistance impacts only spore production. The fitness function is unimodal (left panel). At *t* = 0, the pathogen population is at its mutation-selection equilibrium on the S cultivar (light green distribution). The infection dynamics and phenotypic composition of the pathogen population in the S and R cultivars are displayed in central and right panels, respectively. The black line in the right panel corresponds to the time of emergence *T*_emg_. **Line 2**. Same as line 1 but with *R*_eff_ = 0.99. The fitness function is bimodal with both a global and a local maximum. **Line 3**. The resistance impacts only infection efficiency. The Pairwise Invasibility Plot (PIP) visualizes the sign of the invasion fitness *f*_*x*_(*y*) (left panel). As a mutant strain *y* will invade the resident population *x* if and only if *f*_*x*_(*y*) *>* 0, the PIP reveals a single evolutionary attractor *µ*\* (the vertical line through *µ*\* lies completely inside a region marked ‘−’). **Line 4**. Same as line 3 but with *R*_eff_ = 0.99. The PIP reveals two evolutionary attractors (left panel). For all panels, all other parameters are set to their reference values (Table 2). Note that the time scale axis varies between lines.

In the IE scenario, the pathogenicity trait targeted by the quantitative resistance is only the infection efficiency. The optimization principle based on ℛ_0_ does not hold here, and evolutionary branching and diversification are possible and cannot be solely characterized by the shape of the ℛ_0_. Typically, a polymorphic pathogen population can be selected at equilibrium while ℛ_0_ has a single maximum (Figure S2). In this scenario, we need to calculate the invasion fitness to characterize the evolutionary attractors (equation (3.6)).

As in the SP scenario, the population can evolve to a monomorphic equilibrium (Figure 2 line 3). This issue occurs for a broad choice of resistance genes and deployment strategies (Figure 3). However, a dimorphic population can also be selected (Figure 2 line 4). The dimorphism, easily observed in the airborne pool of spores (Figure S3), is not necessarily so on the individual cultivar (Figure 2 line 4). Indeed, the fitness functions on cultivar can give sharply different relative proportions at equilibrium (Appendix E). A dimorphic population is selected as soon as the cost of adaptation and resistance effectiveness are ≥ 0.9 for most of the proportion of R cultivar at planting except extreme ones (Figure 3, central panel). Dimorphic populations are also selected in a larger number of production situations for the R cultivar characterized by the highest cost of adaptation (*C*_adp_ = 0.99). In this case, the deployment of a proportion of R cultivar above a threshold increasing with decreasing resistance effectiveness leads most of the time to the selection of a dimorphic population (Figure 3, right panel).

**Figure 3:**
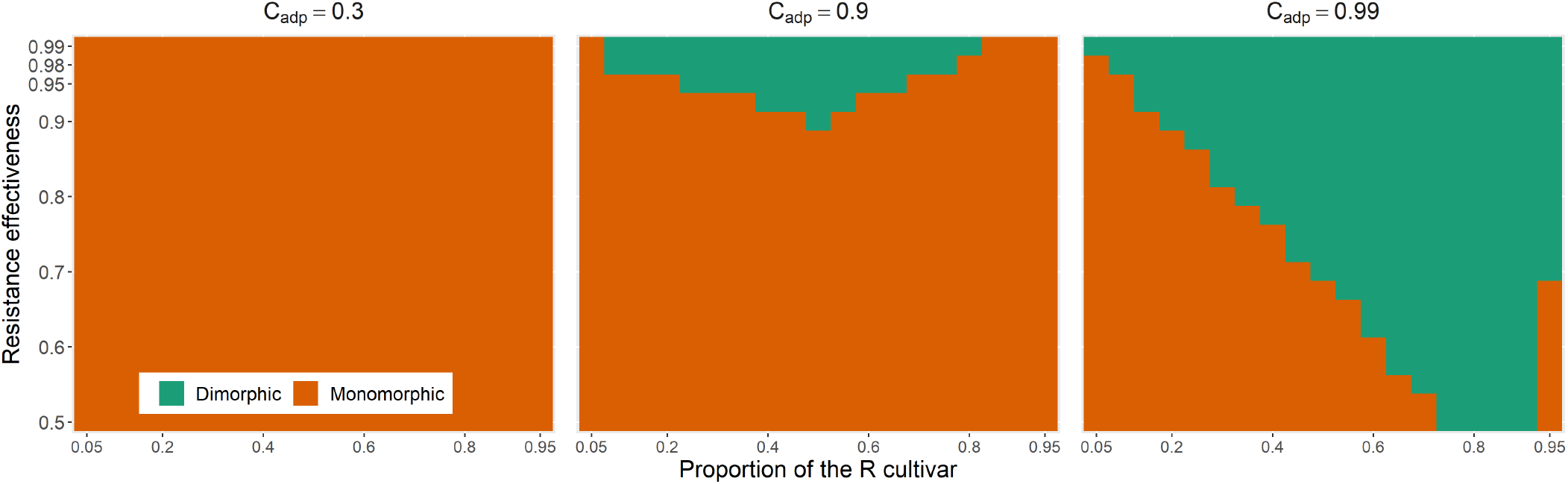
Nature of the evolutionary equilibrium (monomorphic or dimorphic) when the resistance impacts only the infection efficiency. The nature of the equilibrium is characterized as a function of the proportion of the R cultivar at planting (x-axis) and of the relative effectiveness of the resistant cultivar (y-axis) for three costs of adaptation (columns). For all panels, all other parameters are set to their reference values (Table 2).

**Figure 4:**
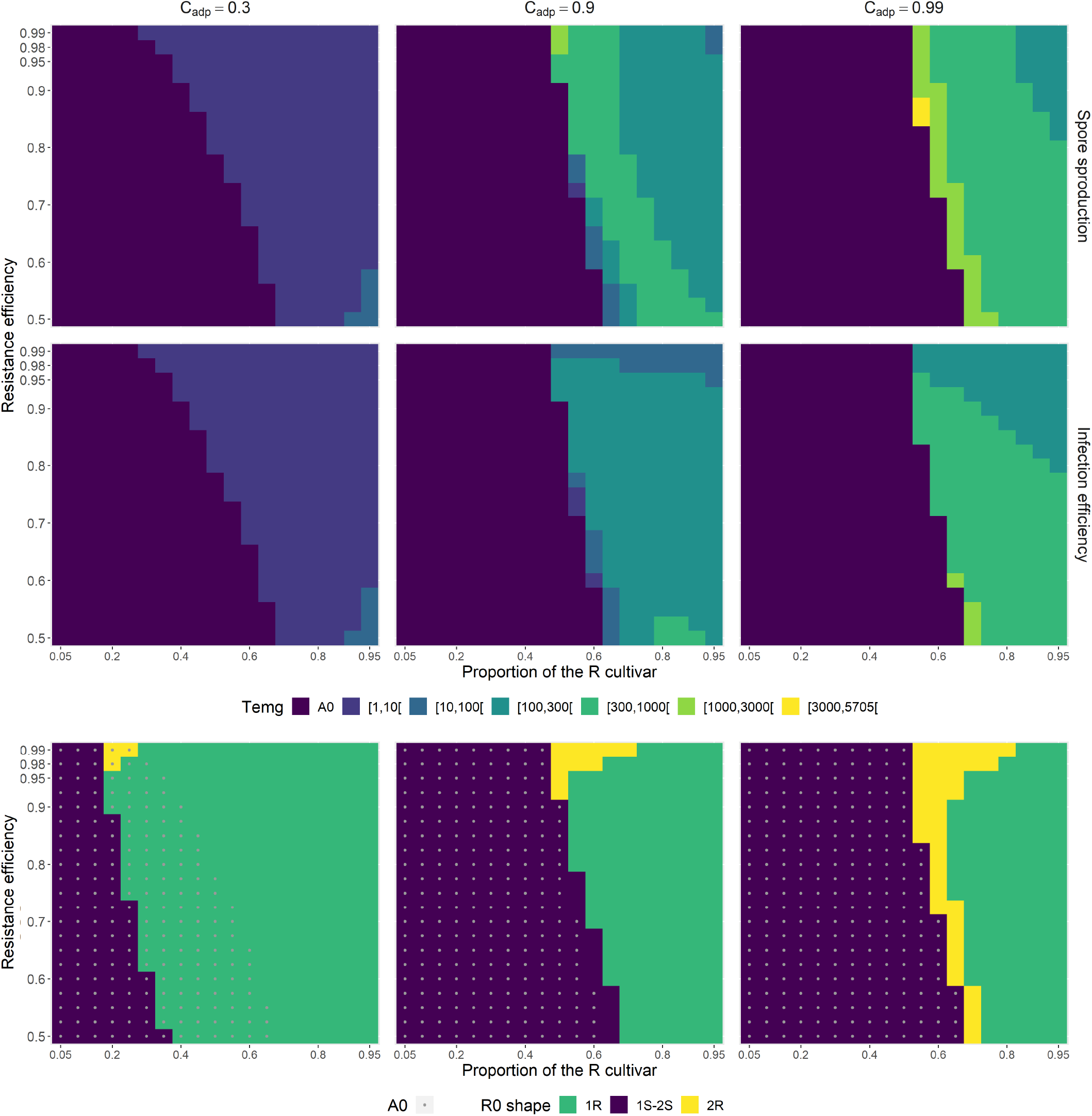
Time to emergence of the evolutionary attractor. The time to emergence *T*_emg_ is the time when the proportion of the evolutionary attractor (the closer to the optimal phenotype on the R cultivar) becomes ≥ 5% in the air pool spores. **Line 1**. The resistance impacts only the total spore production (SP scenario). *T*_emg_ is characterized as a function of the proportion of the R cultivar at planting (x-axis) and of the relative effectiveness of the resistant cultivar (y-axis) for three costs of adaptation (columns). The level A0 indicates situations where the criteria of emergence is meet from the initial conditions. **Line 2**. Same as line 1 when the resistance impacts only the infection efficiency (IE scenario). **Line 3**. Shape of the ℛ_0_ function. Three shapes are classified: (i) “1S-2S” when the evolutionary attractor *µ*\* is closer to *µ*_*S*_ than to *µ*_*R*_ (*i*.*e. µ*\* *<* 0), (ii) “1R” when *µ*\* is closer to *µ*_*R*_ and ℛ_0_ is unimodal, (iii) “2R” when *µ*\* is closer to *µ*_*R*_ and ℛ_0_ is bimodal. The point A0 indicates situations where the criteria of emergence is meet from the initial conditions. For all panels, all other parameters are set to their reference values (Table 2).

The evolutionary epidemiology framework also elucidates the duration of the transient dynamics. Whatever the basic resistance characteristics (pathogenicity trait targeted, resistance effectiveness and cost of adaptation), the relative proportion of the evolutionary attractor is already ≥ 5% in the air pool spores at initial time for a wide range of deployment strategies, mostly as long as the R cultivar is not dominant in the environment (Figure 4, lines 1 and 2, level A0). It mostly corresponds to situations where the evolutionary attractor is closer to *µ*_*S*_ (optimal phenotype on the S cultivar) than to *µ*_*R*_ (optimal phenotype on the R cultivar). In such setting, unless it pre-exists from the beginning in the mutation-selection equilibrium, the evolutionary attractor can be quickly reached by mutations (Figure 2 lines 1 and 3), whether ℛ_0_ has one or two maximums (Figure 4, line 3, level “1S-2S”).

The times to emergence can substantially increase when ℛ_0_ is unimodal but with a maximum getting closer to the optimal phenotype on the R cultivar (Figure 4, line 3, level “1R”). Indeed, more time is then needed to reach the evolutionary attractor by mutation. This increase, small with low cost of adaptation for both scenarios (Figure 4, *C*_adp_ = 0.3), is larger for higher costs (Figure 4, *C*_adp_ = 0.9 or 0.99).

The longest times of emergence are obtained in the SP scenario when ℛ_0_ is bimodal with an global maximum closer to the optimal phenotype on the R cultivar. This configuration is obtained for small ranges of intermediate proportion of R cultivar at planting combined to adaptation cost and resistance effectiveness ≥ 0.9, or combined to the highest adaptation cost considered (*C*_adp_ = 0.99) for all resistance effectiveness tested (Figure 4 line 3, level “2R”). In this configuration, the pathogen population lives for a relatively long time around the initially dominant phenotype and then shifts by mutation to the evolutionary attractor after crossing a fitness minimum. These dynamics occur simultaneously on the S and R cultivars (Figure 2 line 2). The shape of the response surfaces of the times of emergence in the IE scenario are broadly similar to that of the SP scenario. However, in all the production situations explored, the times to emergence are always equal or shorter in the IE scenario than in the SP scenario, notably when the maximum of ℛ_0_ is closer to *µ*_*R*_. Emergence times are shorter by an average of 41 generations when ℛ_0_ is unimodal and by an average of 801 generations when ℛ_0_ is bimodal (Figure 4 line 3, levels “1R” and “2R”). Moreover, in the IE scenario, the evolutionary dynamics differ between cultivars (Figure 2 line 4).

### 4.3 Epidemiological outcomes

The long-term epidemiological control is assessed by the relative Healthy Area Duration gain. In the SP scenario, it ranges from 1.25 to 3.21 with a mean of 1.61 over the whole parameter space explored (Figure 5 line 1). Achieving substantial relative yield gain ≥ 1.5 requires deploying R cultivar with cost of adaptation ≥ 0.9 in at least 50% of the fields regardless of the value of resistance effectiveness. Achieving higher relative yield gain ≥ 2 requires additionally high resistance effectiveness ≥ 0.9.

**Figure 5:**
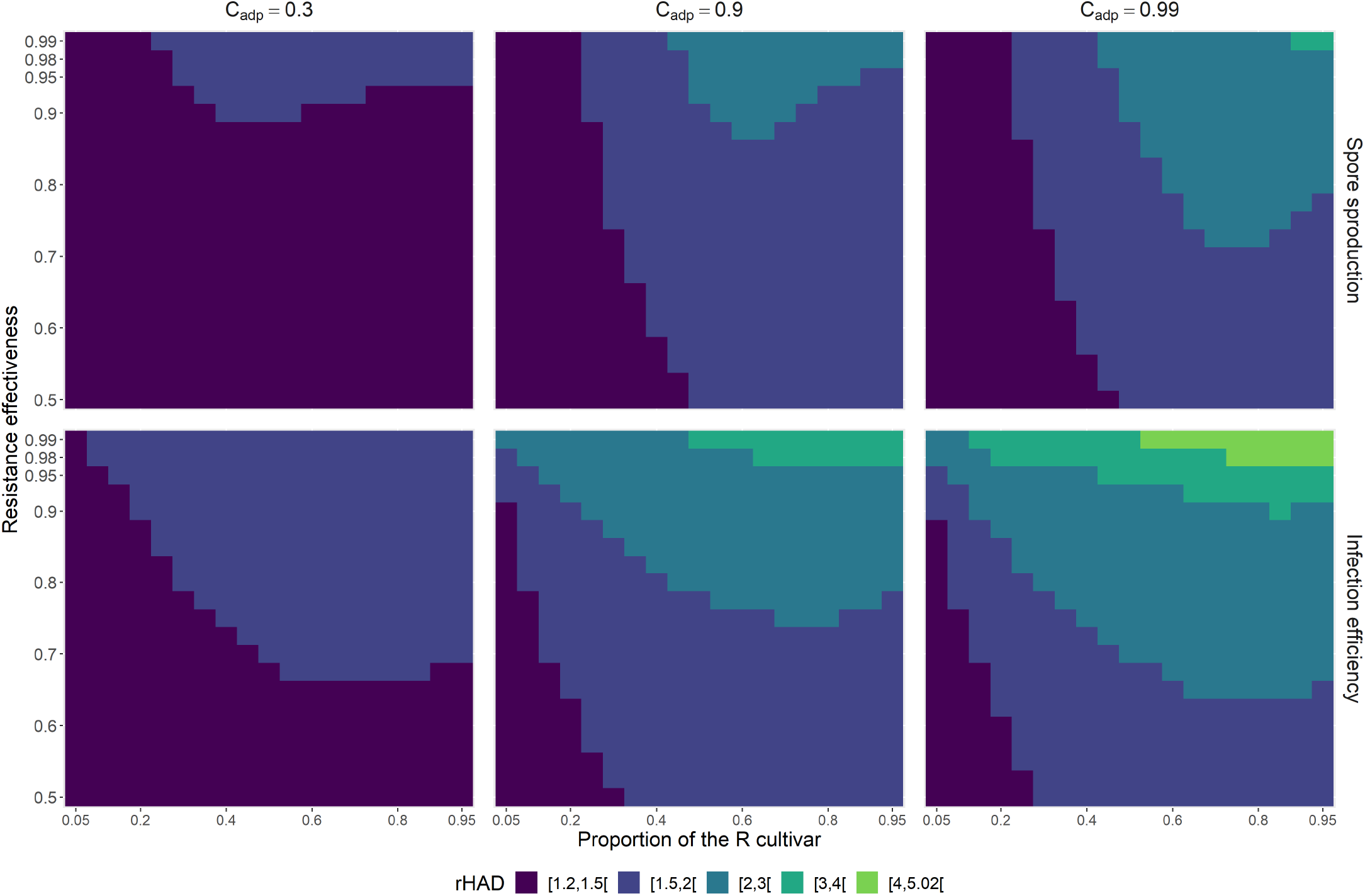
Epidemiological outcomes. The epidemiological control is estimated by the relative Healthy Area Duration (rHAD) gain, a proxy for crop yield. **Line 1**. The resistance impacts only the total spore production (SP scenario). The epidemiological control is characterized as a function of the proportion of the R cultivar at planting (x-axis) and of the relative effectiveness of the resistant cultivar (y-axis) for three costs of adaptation (columns). **Line 2**. Same as line 1 when the resistance impacts only the infection efficiency (IE scenario). For all panels, all other parameters are set to their reference values (Table 2).

In the IE scenario, the relative yield gains range from 1.27 to 5.02 with a mean of 1.9 (Figure 5 line 1). In all the production situations explored, the relative yield gains are higher in the IE scenario than in the SP scenario. The mean (resp. maximal) differences in relative yield gains between them is 0.28 (resp. 1.93). Substantial gains ≥ 2 are obtained for larger ranges of resistance effectiveness and proportions of the R cultivar at planting as soon as costs of adaptation are ≥ 0.9. Remarkably, for both scenarios, the epidemiological control is not correlated to the time to emergence in the production situations explored (compare Figures 4 and 5).

## 5 Discussion

This work follows an ongoing trend aiming to jointly model the epidemiological and evolutionary dynamics of host-parasite interactions. Our theoretical framework, motivated by fungal infections in plants, allows us to tackle the question of the durability of plant quantitative resistances altering specific pathogen life-history traits. In our case study, we show that the evolutionary and epidemiological consequences of deploying resistant cultivars are not necessarily aligned, and depend the pathogen pathogenicity trait targeted by the plant resistance genes. From an evolutionary perspective, the emergence time of the strategy best adapted to the R cultivar tends to be shorter when the resistance impacts infection efficiency (IE scenario) than when it impacts sporulation (SP scenario). In both cases, the emergence time is maximal for an intermediate proportion of R cultivars, as already found by Papa ïx et *al*. (2018). In contrast, from an epidemiological perspective, the epidemiological control is always higher in the IE scenario than in the SP scenario, as already observed by Iacono et *al*. (2012); Rimbaud et *al*. (2018). These general rules are shared among the theoretical studies that have investigated, over the last decade, the epidemiological and evolutionary effects of deploying quantitative resistances in agro-ecosystems, which supports the robustness of these conclusions.

### Bridging the gap between modelling approaches

Although these previous studies all account for the interplay between epidemiological and evolutionary dynamics, they rely on different modeling frameworks, hypotheses and parametrisations. A strong point of our general modelling framework is that it allows us to bridge the gap between different modelling traditions. First, we are able to investigate both the short- and long-term epidemiological and evolutionary dynamics of the host-pathogen interaction (Day & Proulx, 2004). Although the short-term dynamics are investigated numerically, the long-term analysis is analytically tractable and allows us to predict the outcome of pathogen evolution. In contrast to most studies in evolutionary epidemiology, our results are neither restricted to rare mutations, as in the classical adaptive dynamics approach, nor to a Gaussian mutation kernel (see also Mirrahimi (2017)). When the trait distribution is sufficiently narrow, we expect the population to concentrate around the attractors predicted by adaptive dynamics. However, the full model is required to quantify the speed of evolution, and notably the time of emergence of an adapted pathogen strain, which is often more important for practical purposes, than the potential long-term outcome. We also note that “fat-tailed” kernels which allow long-distance dispersal events into the phenotype space could also be considered (Appendix B).

Second, our analysis allows us to consider multimodal fitness functions, which are reminiscent of fitness landscapes in population genetics, and to characterize evolutionary attractors at equilibrium through a detailed description of their shapes (number of modes, steepness and any higher moments with even order). Our results highlight that these features strongly impact transient dynamics which, in turn, shape evolutionary and epidemiological outcomes. In particular, multimodal fitness functions with close local maxima lead to the longest time of emergence of the adapted pathogen strains in the SP scenario. Mathematically, they are characterized by a slow convergence to equilibrium (Burie *al*., 2020). In practice, we could take advantage of these properties through the wise choice of the proportion of each cultivar in the agricultural landscape. Indeed, the shape of the fitness function in this scenario is basically the sum of the fitness in host *k*, weighted by its proportion in the environment (equation 3.3). Third, we propose a simple framework relying on Gaussian distributions to describe the pathogen phenotypic landscape associated with quantitative resistance. Varying the relative effectiveness of resistance and cost of adaptation allows us to describe a whole range of quantitative resistance effects, which allows us to consider the continuum between the two main models used for qualitative plant-parasite interactions: (i) the gene-for-gene model where a parasite strain may have universal infectivity (*R*_eff_ = 1 and *C*_adp_ = 0) and (ii) the matching-allele model where universal infectivity is impossible (*R*_eff_ = 1 and *C*_adp_ = 1) (Thrall et *al*., 2016). Exploring the continuum between these extremes (while noting that *R*_eff_ and *C*_adp_ can only tend to 1 in our framework) allows us to investigate in more detail how the deployment of quantitative resistances impacts epidemiological and evolutionary outcomes.

### Quantitative resistances as a major driver of pathogen evolution

Whether deploying R cultivars in agricultural landscapes will lead to the diversification of an initially monomorphic population and to the long-term coexistence of different pathogen strategies has been investigated using adaptive dynamics by Gudelj et *al*. (2004b) and by Gandon (2004) in the general context of the evolution of virulence of multihost parasites. They highlight that evolutionary outcomes strongly depend on the shape of the trade-off curve between pathogen transmission on sympatric hosts. While concave trade-offs lead to monomorphic evolutionary end-points, convex or sigmoidal trade-offs can lead to evolutionary branching. These results can also explain the existence of sibling fungal pathogens (Gudelj et *al*., 2004a). In their approach, the transmission rate aggregates, in a single proxy trait, infection, spore production and dispersal that jointly impact secondary infection from infected to healthy hosts. By contrast, our framework allows us to disentangle the specific effect of the pathogenicity traits that follow one another during an infection. In agreement with Gudelj et *al*. (2004b), we show that dimorphism is possible with a convex or a sigmoidal trade-off on host infection efficiencies (IE scenario), while the population always remains monomorphic when the trade-off is concave. However, our analysis shows that the predictions crucially depend on which step(s) of the parasite life cycle is (are) affected by quantitative resistance in the R cultivar. The population always remains monomorphic when resistance impacts the sporulation curve only (the total spore production but also any other pathogenicity traits such as the duration of the latency period) irrespective of the underlying trade-off shape (SP scenario). This arises as, in our work, the SP scenario admits an optimization principle based on ℛ_0_ and potential evolutionary attractors are located at the peaks of ℛ_0_ (Lion & Metz, 2018).

Along with the condition leading to their diversification, the evolution of plant pathogens toward generalism or specialism following the deployment of a R cultivar is also a major concern (Papaïx et *al*., 2013, 2014; Croll & McDonald, 2017). We proposed a simple binary criteria classifying the evolutionary attractors as generalist or specialist by comparing their ℛ_0_ on the R and S cultivars to a preset threshold (Figure S4). It highlights that high costs of adaptation are a major factor leading to the selection of specialists but also that emergence times and epidemiological control are weekly structured by this classification (Figure S4). Whether favoring deployment strategies leading to the selection of generalist or specialist pathogen strains is an open question that deserves further investigation. It depends, among other things, on the pre-existing level of pathogen diversification (Papaïx, 2011). However, addressing this question in the context of quantitative resistance firstly requires new indices measuring a degree of generalism within a specialist/generalist continuum.

### ℛ_0_ in heterogeneous host environments sharing a common pool of propagules

Many practical epidemiological studies rely on the concept of the basic reproduction ratio ℛ_0_. For instance, van den Bosch et *al*. (2008) calculate ℛ_0_ for lesion-forming foliar pathogens in a setting with two cultivars but without effect of the age of infection on sporulation and on disease-induced mortality. While ℛ_0_ is typically calculated using the spectral radius of the next generation operator (Diekmann et *al*., 1990), we follow here the methodology based on the generation evolution operator (Inaba, 2012) to derive an expression for the basic reproduction number ℛ_0_ in heterogeneous host populations with any number of cultivars. This expression captures the time-scales inherent to the life-cycle of plant fungi pathogens with, notably, time-varying sporulation. Capturing such patterns is a challenge in modelling plant diseases identified by Cunniffe *al*. (2015). Moreover, with a common pool of well-mixed airborne pathogen propagules, the function ℛ_0_(*x*) is an exact fitness proxy for competing strains differing potentially in their sporulation curve (including the latent period, the total spore production and shape parameters). This deserves to be mentioned as usually the computation of ℛ_0_(*x*) assumes that the invading pathogen enters an uninfected host population. However, more generally, a clear distinction between pathogen invasion fitness *R*(*x, y*) and epidemiological ℛ_0_(*x*) is necessary to properly discuss the adaptive evolution of pathogens (Lion & Metz, 2018). Even with a common pool of spores, the optimization principle of ℛ_0_(*x*) does not hold when infection efficiencies differ between host classes.

### Notes on some model assumptions

The model assumes an infinitely large pathogen population. Demographic stochasticity is thus ignored while it can impact evolutionary dynamics (*e*.*g*., lower probabilities of emergence and fixation of beneficial mutations, and reduction of standing genetic variation (Kimura, 1962)). In particular, genetic drift is more likely to impact the maintenance of a neutral polymorphism rather than of a protected polymorphism where selection favors coexistence of different genotypes against invasions by mutant strategies (Geritz et *al*., 1998). The effect of genetic drift depends on the stability properties of the model considered. As our model has a unique globally stable eco-evolutionary equilibrium, genetic drift is likely to have a weaker impact than in models with several locally stable equilibria. Moreover, the large *N*_*e*_ (from 10^3^ to 3.10^4^) reported at field scale for several species of wind-dispersed, spore-producing plant pathogens (Ali et *al*., 2016; Zhan et *al*., 2001; Walker et *al*., 2017) suggest a weak effect of genetic drift.

The model assumes that the aggressiveness components are mutually independent. However, correlations between traits have sometimes reported. For instance, Pariaud et *al*. (2013) observed a positive correlation between the duration of the latent period and fecundity of a plant fungi. This relationship describes a phenotypic trade-off because a short latent period and a high sporulation probability represent fitness advantages. It can be introduced into the SP scenario assuming that the latent period and the total spore production of a pathogen strain *x* are linked by some mathematical functions, such as the quadratic relationship suggested by Pariaud et *al*. (2013). Alternatively, correlations between the pathogen life-history traits can emerge from the covariance matrix of the (multidimensional) mutation kernel (Gandon, 2004). Another feature of the framework that we did not use is the possibility to track the evolution of the pathogen’s quantitative traits using Price’s equation (Day & Proulx, 2004; Day & Gandon, 2007; Iacono et *al*., 2012). Indeed, differential equations for mean phenotype and phenotypic variance of any trait of interest can be derived from model (2.2).

The model also assumes a unique pool of well-mixed propagules. Thus, spore dispersal disregards the location of healthy and infected hosts. This assumption, which ensures the one-dimensional environmental feedback loop of the model, is more likely when the extent of the field or landscape considered is not too large with respect to the dispersal function of airborne propagules. Airborne fungal spores often disperse over substantial distances. Mean dispersal distances range from 100 m to 1 km and, in most cases, long-distance dispersal events are frequent (Fabre et *al*., 2021). Recently, a spatially implicit framework embedded into integro-differential equations was used to describe the eco-evolutionary dynamics of a phenotypically structured population subject to mutation, selection and migration between two habitats (Mirrahimi, 2017). With the same goal Rimbaud et *al*. (2018) used a spatially-explicit framework with several habitats embedded into a HEIR stochastic model. It would be interesting to draw on these examples and extend our approach to a spatially explicit environment. Indeed, when dispersal decreases with distance, large homogeneous habitats promote diversification, while smaller habitats, favoring migration between distinct patches, hamper diversification (Débarre & Gandon, 2010; Haller et *al*., 2013; Papaïx et *al*., 2013, 2014).

### Applied significance

Three applied messages deserve to be mention. The first one is the equation of ℛ_0_ in an environment with several cultivars. This equation relies on a gamma sporulation curve documented for several plant fungi (van den Bosch et *al*., 1988; Sache et *al*., 1997; Kolnaar & Bosch, 2011; van den Bosch et *al*., 1988; Segarra et *al*., 2001) and explicitly integrates the pathogenicity traits expressed during the basic infection steps of a plant fungi (infection efficiency, latent period, sporulation dynamics). As these traits can be measured in the laboratory, this expression of ℛ_0_ bridges the gap between plant-scale and epidemiological studies, and between experimental and theoretical approaches. It helps address the practical difficulty of estimating _0_ for real populations pointed out by Gilligan and van den Bosch (2008). For example, it can be used to compare the fitness of a collection of pathogen isolates (*e*.*g*. see Montarry et *al*. (2010) for an application to potato late blight) or to predict their field structure (*e*.*g*. see Durand et *al*. (2017) for an application to Lyme disease).

The second applied message is that the choice of quantitative resistance genes operated by plant breeders is a driver of pathogen diversification mediated by the parasite life cycle targeted by the resistance. Quantitative resistances specifically impacting distinct stages of pathogen cycles have been identified in several pathosystems (*e*.*g*. Azzimonti et *al*. (2013); Chung *al*. (2010); Jorge et *al*. (2005)). Moreover, understanding the conditions that maintain pathogen polymorphism is a long-standing question in disease ecology and evolution, with implications for disease management (Vale, 2013). Here we show that the pathogen population always remains monomorphic when resistance impacts sporulation only irrespective of the underlying trade-off shape. In contrast, pathogen diversification is possible when resistance impacts infection efficiency only. It highlight that a detailed knowledge of the effect of resistance genes on pathogen life-cycles is necessary to fully understand how the deployment of quantitative resistances will impact strain structures underlying the patterns of disease incidence. This is an important message as quantitative resistances are now increasingly used by plant breeders in response to the rapidity with which major resistance genes are often breakdown (Zhan et *al*., 2015). Interestingly, similar questions arise in the literature on the epidemiological and evolutionary consequences of vaccination strategies. For instance, quantitative resistance traits against pathogen infection rate, latent period, and sporulation rate are analogous to partially effective (imperfect) vaccines with anti-infection, anti-growth or anti-transmission modes of action, respectively (Gandon et *al*., 2001). Similarly, the proportion of R cultivar deployed is analogous to the vaccination coverage in the population; the relative effectiveness of resistance is termed vaccine efficacy and the relative cost of adaptation is termed cost of escape (Gandon & Day, 2007).

Finally, we confirm, following Papaïx et *al*. (2018) but with a different theoretical approach, that “there is no silver bullet deployment strategy” maximising at the same time epidemiological and evolutionary outcomes (Rimbaud et *al*., 2021). Accordingly, as the different stakeholders involved in plant resistance management may pursue objectives not always compatible (e.g., growers, breeders), they must all be involved to define the best management strategies.

## Code availability

The MATLAB codes used to simulate the model and generate the main figures have been deposited in Dataverse at https://doi.org/10.15454/WAEIMA

## A Supplementary figure

**Figure S1:**
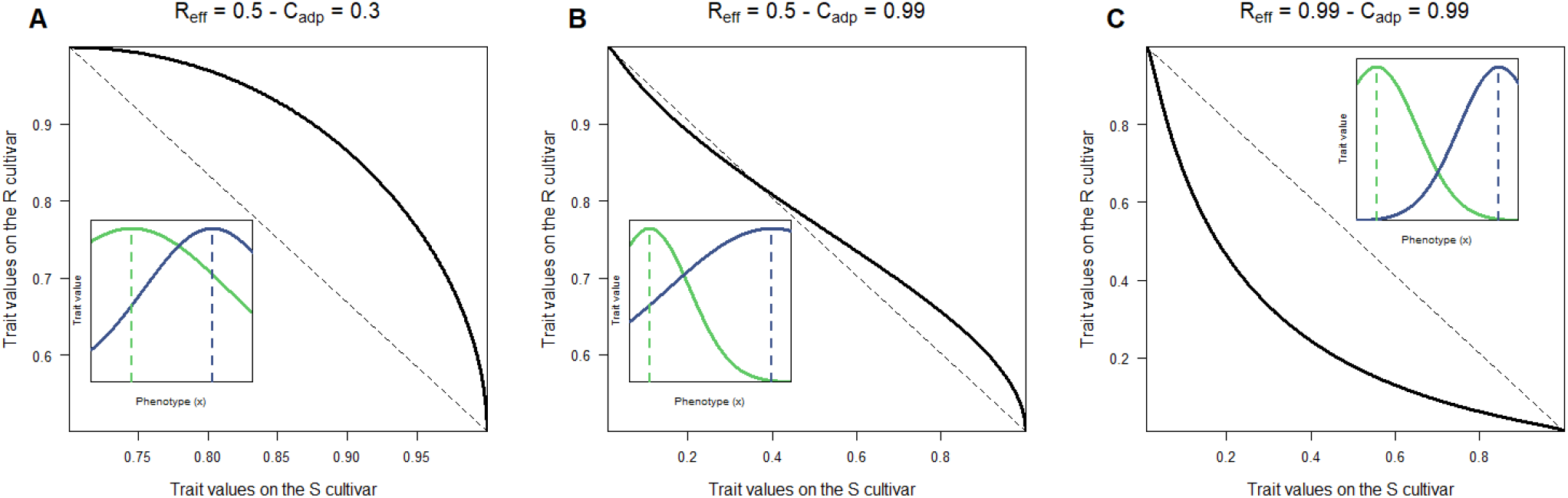
Shapes of the trade-off curves of the pathogen traits values on the S cultivar against those on the R cultivar as given by *ω*_*R*_ = *f* (*ω*_*S*_) with *x* ∈ [*µ*_*S*_, *µ*_*R*_]. The trade-off can be either concave (panel A), sigmoidal (panel B) or convexe (panel C). Inset graphs show the corresponding Gaussian functions *ω*_*S*_(*x*) and *ω*_*R*_(*x*) as in Figure 1B. For all panels all other parameters are set to their reference values (Table 2).

**Figure S2:**
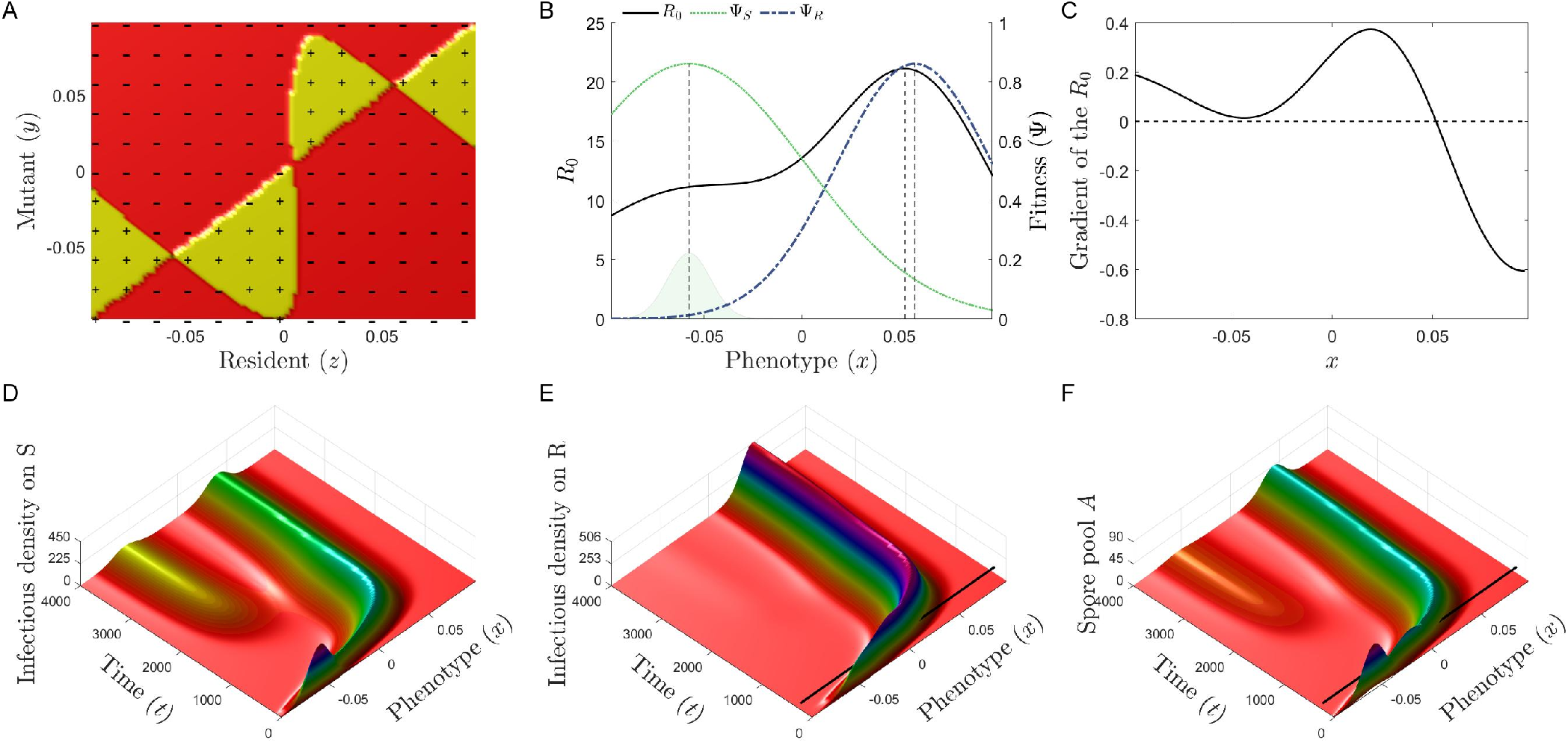
An example of configuration where a polymorphic pathogen population is selected at equilibrium (panels A, D, E, F) while a single local maximum exists for *R*_0_ (panel B) as confirmed by its gradient (panel C). Parameters values are *ϕ* = 0.64, *σ*_*S*_ = 0.06, *C*_adp_ = 0.845 and *R*_eff_ = 0.985 and *ε* = 0.002. Other parameters are set to their reference values. The PIP in panel A visualizes the sign of the invasion fitness *f*_*x*_(*y*). The mutant strain *y* will invade the resident population *x* if and only if *f*_*x*_(*y*) *>* 0.

**Figure S3:**
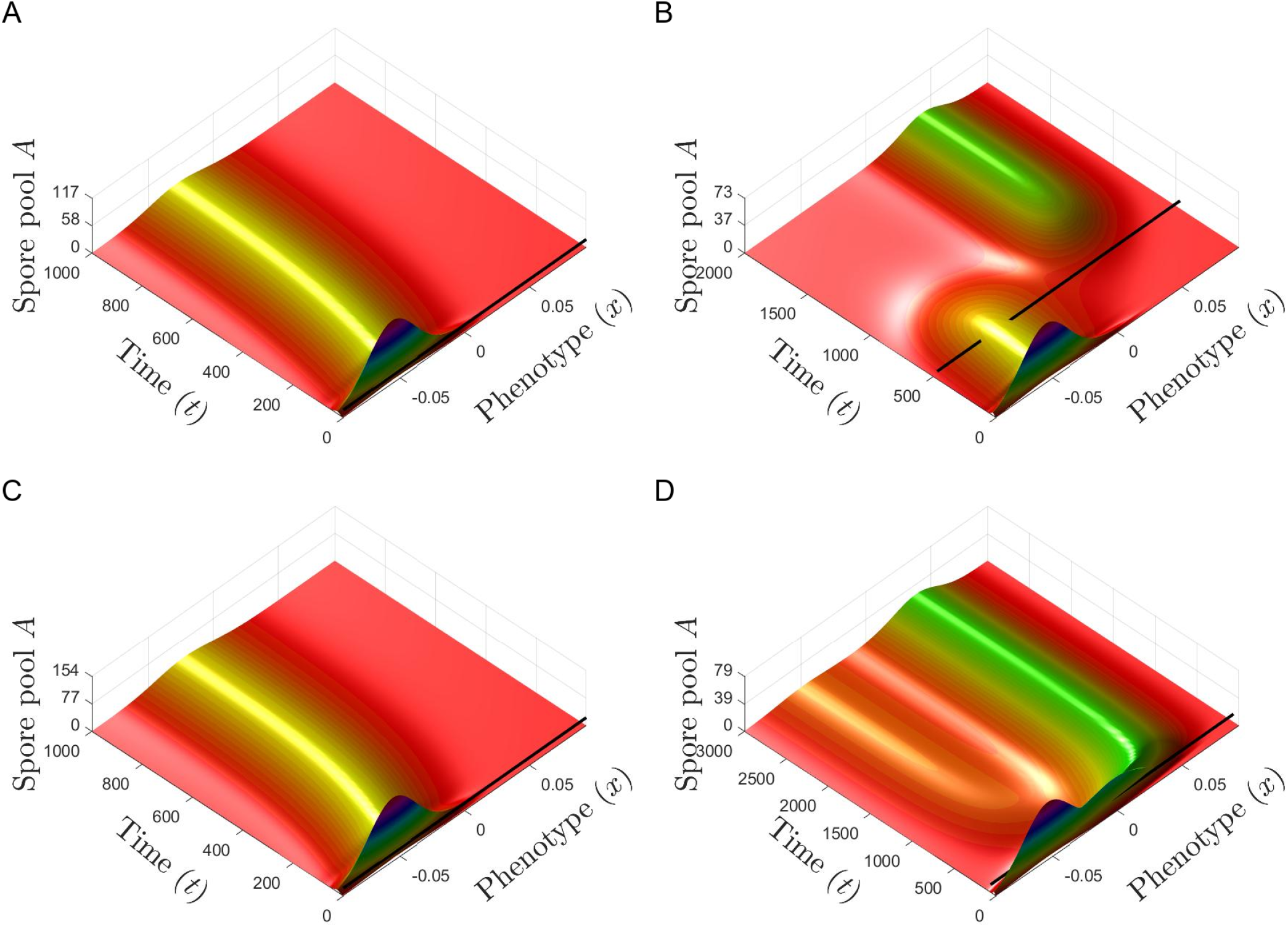
Dynamics of the density of airborne pool of pathogen *A*(*t, x*). **A-B**. The resistance impacts the total spore production (SP scenario). The dynamics of *A*(*t, x*) corresponds to the lines 1 and 2 of Figure 2: (A) *ϕ* = 0.5, *C*_adp_ = 0.8 and *R*_eff_ = 0.5 and (B) *ϕ* = 0.5, *C*_adp_ = 0.8 and *R*_eff_ = 0.99. **C-D**. The resistance impacts the pathogen infection efficiency (IE scenario). The dynamics of *A*(*t, x*) corresponds to the lines 3 and 4 of Figure 2: (C) *ϕ* = 0.5, *C*_adp_ = 0.8 and *R*_eff_ = 0.5 and (D) *ϕ* = 0.5, *C*_adp_ = 0.8 and *R*_eff_ = 0.99. For all panels, all other parameters are set to their reference values (Table 2).

**Figure S4:**
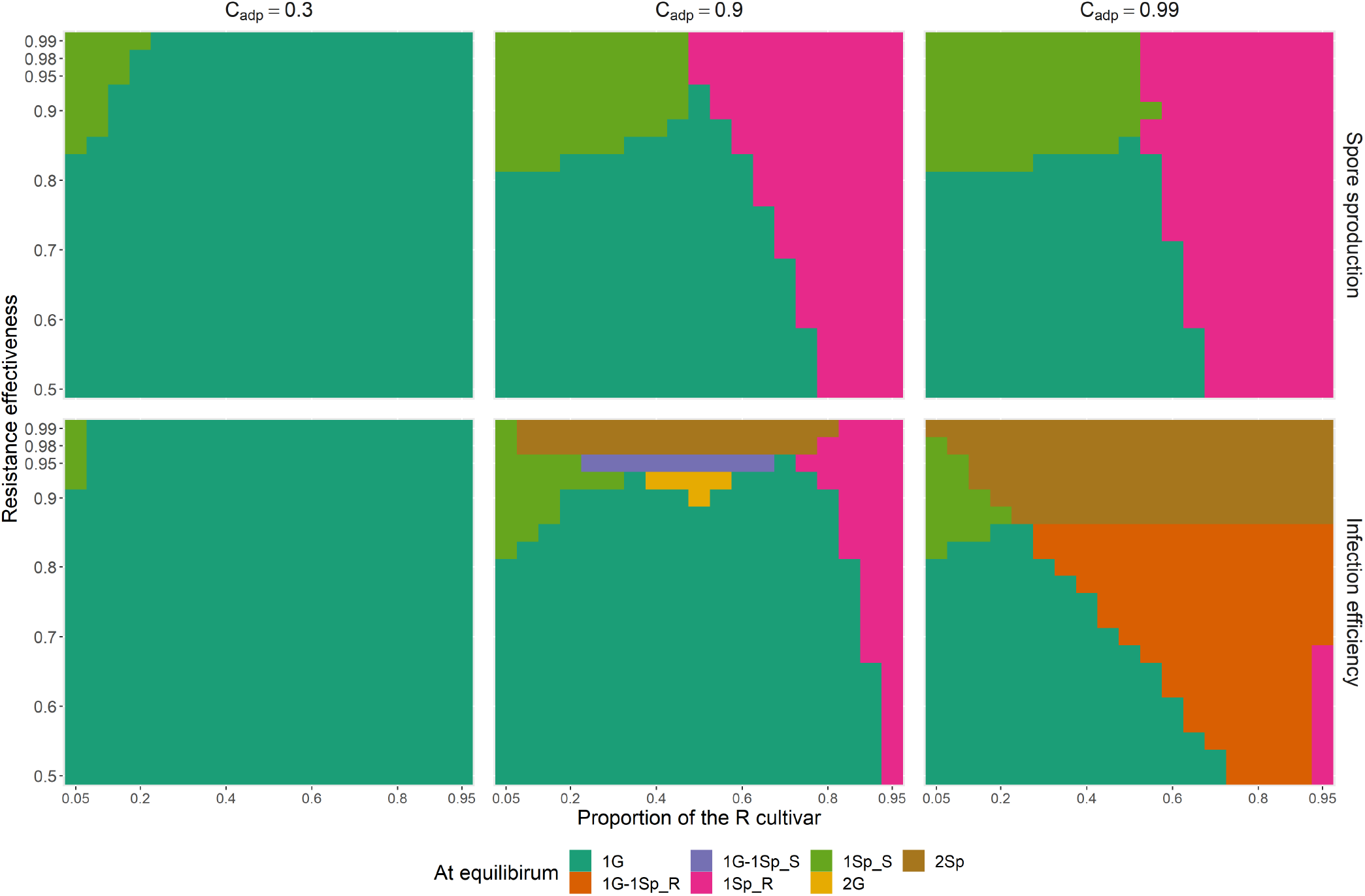
Characterization as generalist and/or specialist of the evolutionary attractors. Due to the quantitative interactions considered, a phenotype *µ*\* has never strictly a single host range, and its characterization depends on a threshold set here to 0.2. Accordingly, *µ*\* is termed a specialist of the R cultivar if 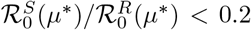, a specialist of the S cultivar if 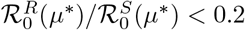, and a generalist otherwise. **Line 1**. The resistance impacts only the total spore production (SP scenario). The evolutionary attractor is characterized as a function of the proportion of the R cultivar at planting (x-axis) and of the relative effectiveness of the resistant cultivar (y-axis) for three costs of adaptation (columns). The equilibrium is always monomorphic in the SP scenario with one generalist (1 G), one specialist of the S cultivar (1 SpS) or one specialist of the R cultivar (1 SpR). **Line 2**. The resistance impacts only the pathogen infection efficiency (IE scenario). In the IE scenario, the equilibrium can also be dimorphic with two specialists (2 Sp), one generalist and one specialist of the S cultivar (1G & 1 SpS), one generalist and one specialist of the R cultivar (1G & 1 SpR) or two generalists (2 G). For all panels, all other parameters are set to their reference values (Table 2).

## B Properties of the mutation kernel *m*_*ε*_

In the model, mutations randomly displace strains into the phenotype space at each generation according to the kernel *m*_*ε*_. In the simulations, we used a centered multivariate Gaussian distribution with standard deviation *ε*. It leads to 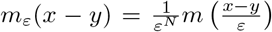. However, the kernel is not restricted to Gaussian distributions. For example, any exponential-power kernels are possible and, in particular, “fat-tailed” exponential-power kernels which allow long-distance dispersal events into the phenotype space can be considered (Klein et *al*., 2006).

More generally, the kernel function *m*_*ε*_ arising in model (2.2) should satisfy the following properties:

**(H1)** The function *m*_*ε*_ is almost everywhere strictly positive on ℝ_*N*_ and should be normalised such that,

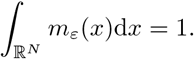

This last condition expresses that all interactions generated on the phenotypic space of pathogens necessarily end up somewhere on that space.

**(H2)** Its variation should only depend on the distance separating the points between which the interactions are evaluated (*i*.*e. m*_*ε*_(*x*) = *m*_*ε*_(−*x*), for all *x* ∈ ℝ_*N*_).

**(H3)** It decays rather fast at infinity in the sens that *m*_*ε*_(*x*) = *ε*^−*N*^ *m*(*x/ε*) and 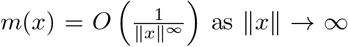as ‖*x*‖ → ∞. In other words, 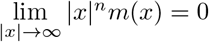, for all *n* ∈ ℕ. This assumption does not mean the mutation kernel has a very fast decay at infinity. We emphasize that the decay of the mutation kernel distribution considered here allows considering the tails of a wide variety of distributions.

## C Some special cases of the general model (2.2)

By omitting the age structure, we re-write model (2.2) as follows

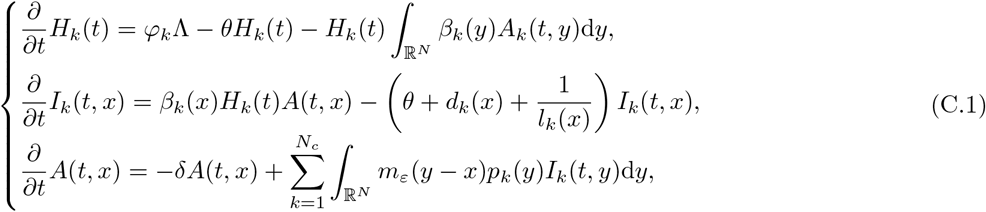

wherein we take into account the (host and strain-specific) duration of the sporulation period, denoted by *l*_*k*_(*x*).

Furthermore, if we assume that there are no “interactions” in the phenotypic space of pathogens, *i*.*e*. without mutations: *ε* → 0, then the simplified model (C.1) rewrites

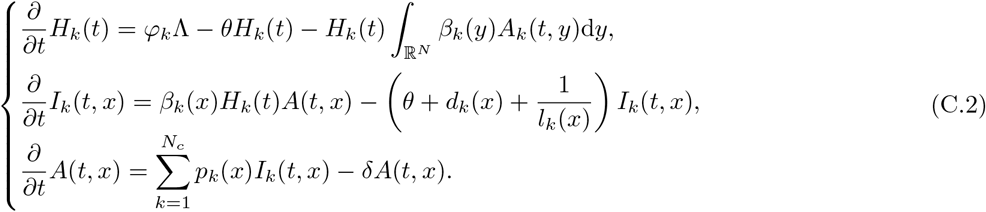

## D The fitness function

In this appendix we explain how to compute the fitness function. To that aim, by formally taking the limit *ε* → 0 into (2.2), this system becomes

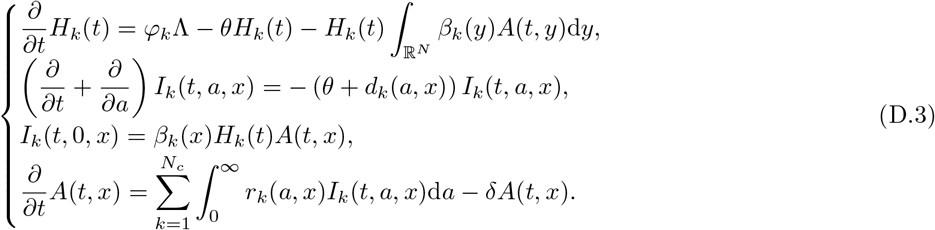

Let us assume that system (D.3) reaches a monomorphic epidemiological equilibrium 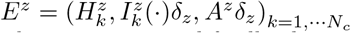, for some trait *z*, before a new mutation with trait value, say, *y* occurs. Note that *E*^z^ is the environmental feedback of the resident *z*. We introduce a small perturbation in (D.3) in the phenotype trait *y*, so that the evolution of the system reads as follows: 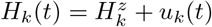 and

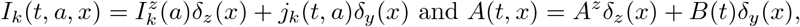

and the small perturbations for the infection, *j*_*k*_ and *B*, are governed by the linearized system of equations around *E*^*z*^. This reads as

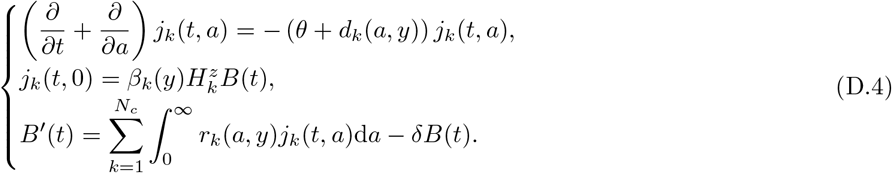

In order to study the evolution of this perturbation we will derive a renewal equation on *b*^*z*^(*t, y*), the density of newly produced spores at time *t* with phenotype *y* in the resident population with phenotype *x*. This term is more precisely defined by

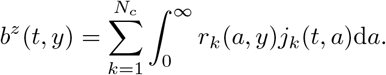

It then follows from the *j*_*k*_-equation of the linear system (D.4), that

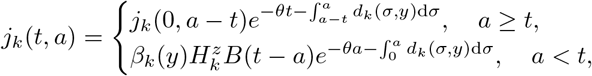

while

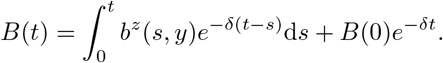

As a consequence, *b*^*z*^(*t, y*) satisfies the following renewal equation:

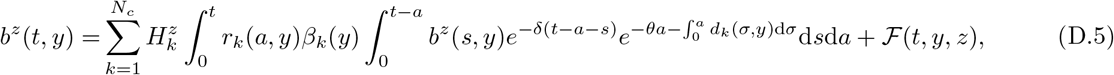

wherein we have set

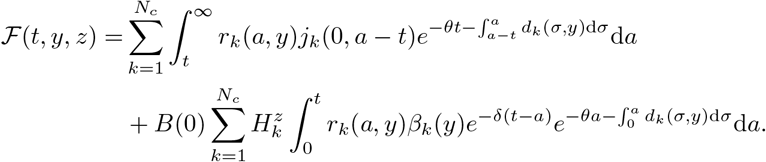

Then (D.5) can be rewritten as

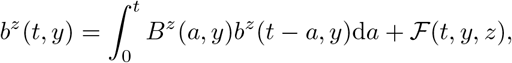

where *B*^*z*^(*a, y*) is the expected number of new infections produced per unit time, in a resident host population with phenotype *z*, by an individual which was infected *a* units of time ago with the phenotype *y*, given by

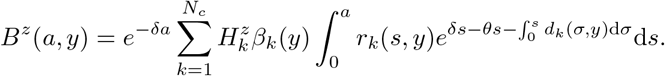

Due to the above formulation, it follows from classical adaptive dynamics (Diekmann et *al*., 2005; Geritz et *al*., 1997; Metz et *al*., 1996) that the spore numbers, ℛ (*y, E*^*z*^), of a rare mutant strategy, *y*, in the resident *z*-population is given by

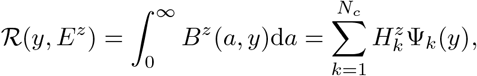

wherein 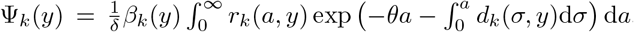. Then, the invasion fitness *f*_*z*_ (*y*) of a mutant strategy *y* in the resident *z*-population is given by

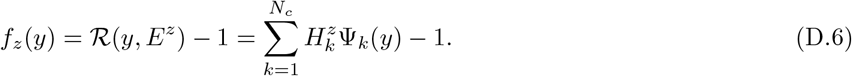

Note that when the environmental feedback *E*^*z*^ is reduced to the disease-free environment, then 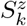 re-writes as 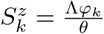. And the epidemiological basic reproduction number of the pathogen with the phenotype *y* is calculated as

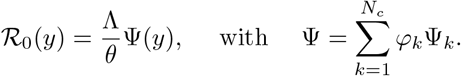

Once the pathogen has spread and reached the monomorphic equilibrium, the endemic feedback environment *E*^*z*^ becomes

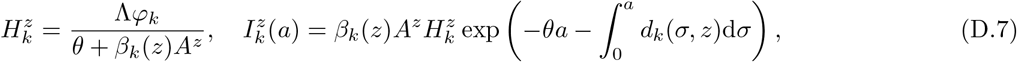

where *A*^*z*^ *>* 0 is the unique solution of the following equation (only defined when *R*_0_(*z*) *>* 1):

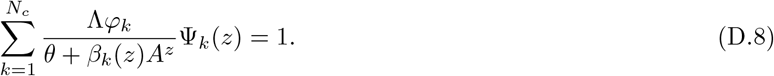

## E Dimorphic or monomorphic equilibrium

To simplify the presentation, we consider system (2.2) with *N*_*c*_ = 2 corresponding to S and R cultivars. Denote by (*H*_0_, *i*_0_(·), *A*_0_) the endemic equilibrium of system (2.2) as *ε* → 0 and when only S is cultivated (*i*.*e*. when the proportion *φ* of R is zero). From results in Djidjou-Demasse et *al*. (2017) we have

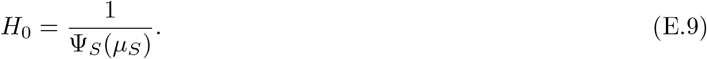

Now, let (*H*_*S*_, *H*_*R*_, *i*_*S*_(°), *i*_*R*_(°), *A*) be an equilibrium of system (2.2) when a proportion *φ >* 0 of R is cultivated Next recall that, for *k* ∈ *{S, R}*,

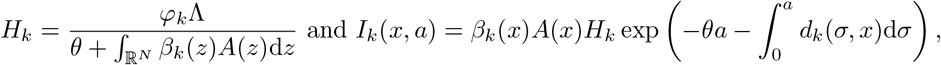

so that *A*(·) becomes a solution of the nonlinear equation:

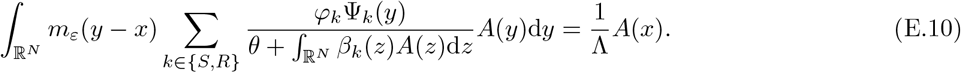

Using this equation we heuristically explore conditions yielding to dimorphic or monomorphic equilibrium.

### Quantitative resistance impacting infection efficiencies *β*_*S*_ and *β*_*R*_ (IE scenario)

We formally assume that the population of spores writes 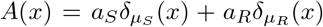, and we plug this ansatz into equation (E.10) above. This yields, for any *x*,

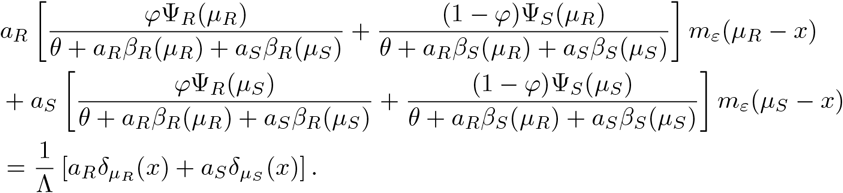

Letting *ε* → 0 and recalling that *m*_*ε*_(*x*) ≈ *δ*_0_(*x*), one obtains

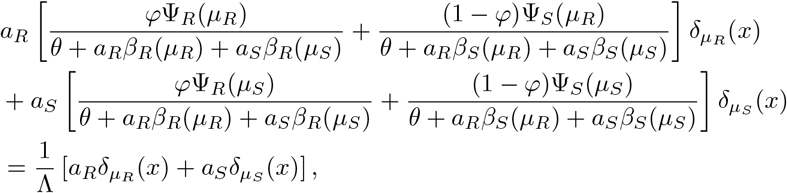

that is

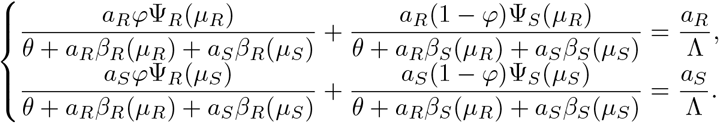

As a consequence, for the equilibrium to be dimorphic, namely *a*_*R*_ *>* 0 and *a*_*S*_ *>* 0, it is necessary that there exist *a*_*R*_ *>* 0 and *a*_*S*_ *>* 0 satisfying the following system of equations:

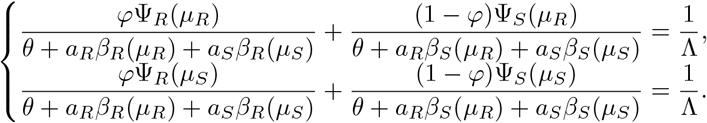

We set

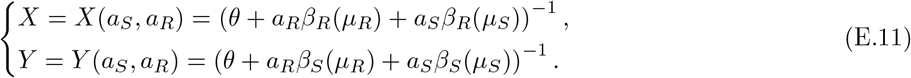

Recall that with the IE scenario we have *r*_*S*_ = *r*_*R*_ and *d*_*S*_ = *d*_*R*_ such that the fitness function

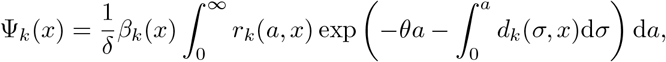

takes the form Ψ_*k*_ = *c*_0_*β*_*k*_ for *k* = *S, R* (where *c*_0_ is the same positive funtional for S and R). Doing that, the above system rewrites

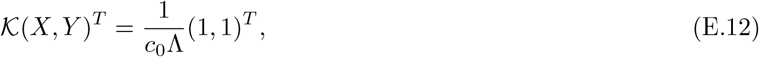

wherein 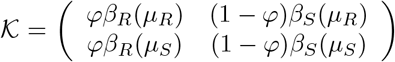 and *c*_0_ = *c*_0_(*µ*_S_) = *c*_0_(*µ*_R_). With the IE scenario (*i*.*e*. with trade-off on infection efficiency *β*_*k*_), we reasonably have *β*_*R*_(*µ*_*R*_) *> β*_*R*_(*µ*_*S*_) and *β*_*S*_(*µ*_*S*_) *> β*_*S*_(*µ*_*R*_). Therefore, det(𝒦) = *φ* (1 − *φ*) (*β*_*R*_(*µ*_*R*_)*β*_*S*_(*µ*_*S*_) − *β*_*R*_(*µ*_*S*_)*β*_*S*_(*µ*_*R*_)) *>* 0. Then, solving system (E.12) for (*X, Y*) yields to

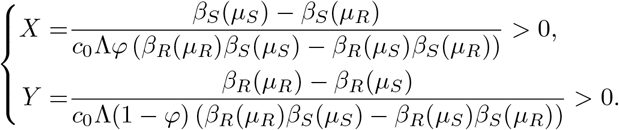

Since 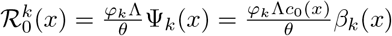, the above system rewrites

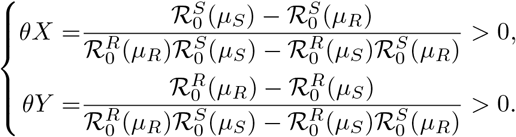

Coming back to the definition of *X* = *X*(*a*_*S*_, *a*_*R*_) and *Y* = *Y* (*a*_*S*_, *a*_*R*_) provided by (E.11), we then find

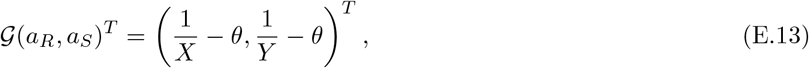

with 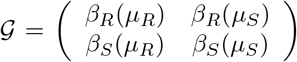. Because det(𝒢) = (*β*_*R*_ (*µ*_*R*_)*β* _*R*_ (*µ*_*R*_) *β*_*R*_ (*µ*_*R*_)*β*_*R*_ (*µ*_*R*_)) *>* 0, it comes that for the equilibrium to be dimorphic it is necessary that

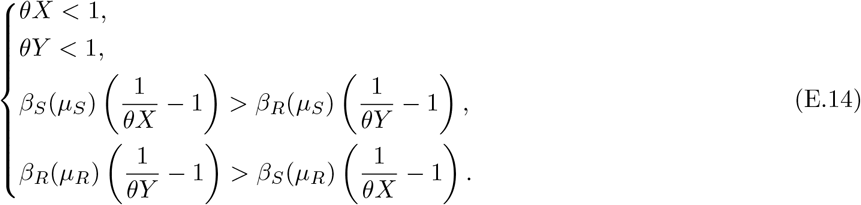

This heuristic condition (E.14) is necessary (but not sufficient) for system (2.2) (here with *N*_*c*_ = 2) to admit an endemic dimorphic equilibrium. The situation with a technical assumption on disjoint supports of *β*_*k*_, is rigorously studied in Burie *al*. (2019).

But here, in order to go slightly further in our analysis, we assume a strong trade-off on infection efficiency, namely

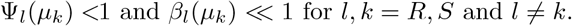

We deduce that the above system of equation roughly simplifies into

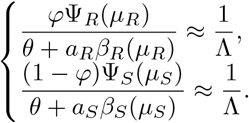

Hence the proportions of each phenotype, *µ*_*S*_ and *µ*_*R*_, can be calculated as

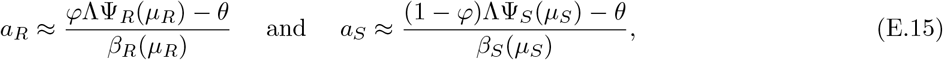

provided the following threshold conditions in this strong trade-off framework

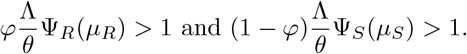

### Quantitative resistance impacting total sporulation production *p*_*S*_ and *p*_*R*_ (SP scenario)

In this case, using the same argument as in Djidjou-Demasse et *al*. (2017) we can prove that the spore population is monomorphic at equilibrium such that *A*(*x*) = *a*\**δ*_*µ*_* (*x*); with *a*\* *>* 0, providing that we are not in a strict symmetric configuration of the fitness function. Applying the same arguments as in the previous section leads to

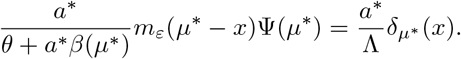

Again with *ε* → 0, it comes

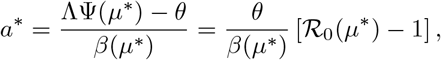

with *R*_0_(*µ*\*) *>* 1.

### F ℛ_0_ as the fitness proxy

By Equations (D.6) and (D.7), it comes

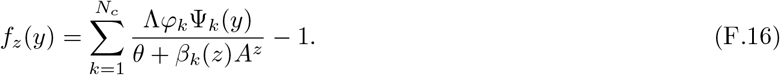

Using Equation (D.8) defining the resident equilibrium, i.e. *R*(*z, E*^*z*^) = 1, (F.16) becomes

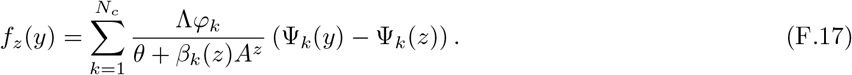

**When infection efficiencies do not differ between host classes (***i*.*e. β*_*k*_ = *β*(*x*), **for every** *k* **and every** *x*). Then (F.17) gives

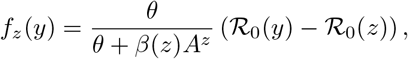

and then

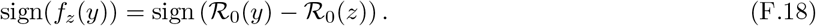

